# 23S rRNA modifications stimulate catalytic activity and prevent the formation of alternative structures

**DOI:** 10.64898/2026.03.01.708810

**Authors:** Daniel S. D. Larsson, Aivar Liiv, Rya Ero, Jaanus Remme, Maria Selmer

**Author notes:** Daniel S.D. Larsson and Aivar Liiv contributed equally to this work.

## Abstract

Ribosomal RNA modifications cluster around the peptidyl transferase center (PTC), the catalytic center of the ribosome, yet their collective functional roles remain unclear. Here we analyze *Escherichia coli* ribosomes lacking 11 or 12 modifications near the PTC. Using kinetic assays, we show these hypo-modified ribosomes catalyze peptide bond formation at rates two- to threefold lower than wild-type and exhibit reduced thermal stability. Cryo-electron microscopy of hypo-modified ribosomes reveals multiple alternative conformations of the PTC and exit tunnel regions, disrupting native stacking and hydrogen bonding critical for positioning of tRNA substrates. These findings indicate that rRNA modifications stabilize the native PTC structure, preventing formation of alternative, nonfunctional conformations and thereby enhance catalytic efficiency. Our study provides insight into how rRNA modifications fine-tune ribosome function by maintaining structural integrity essential for efficient translation

**GRAPHICAL ABSTRACT:** 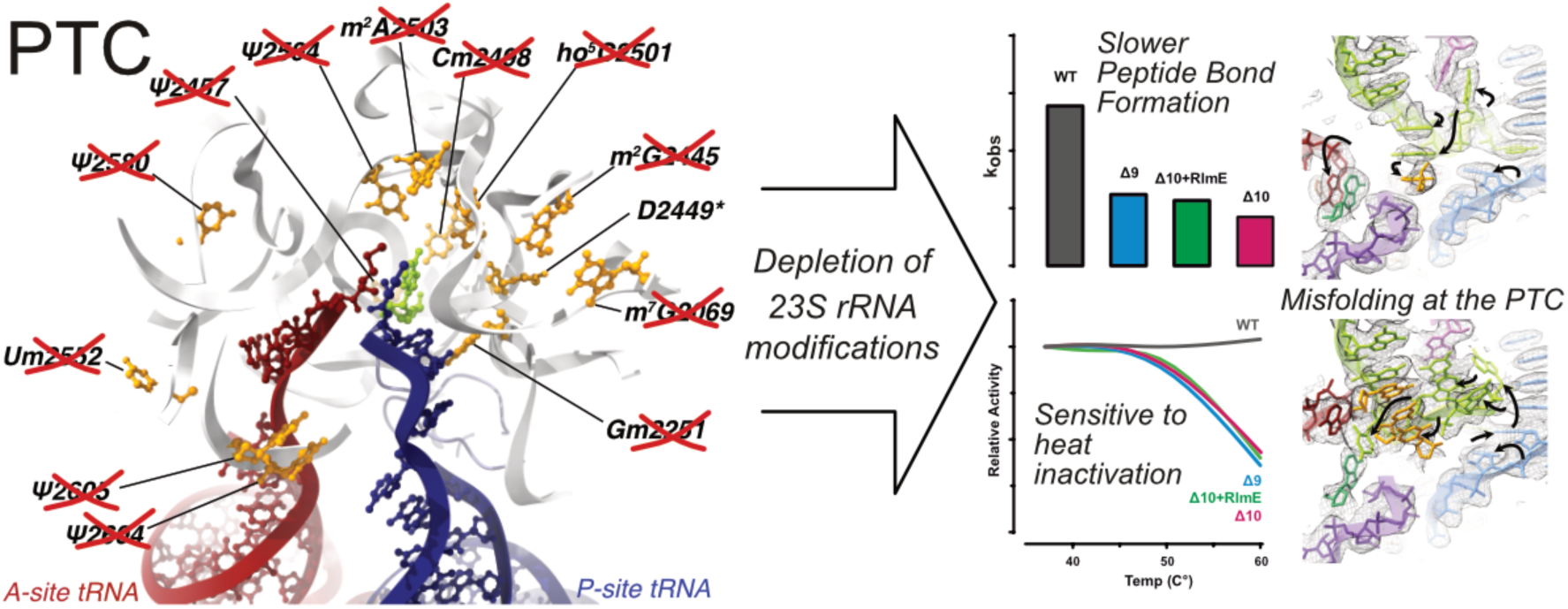

## INTRODUCTION

Stable RNA species contain modified nucleotides that are often located at conserved sites. In the ribosomal RNA (rRNA) of bacteria, most modified nucleotides (MNs) are located around the decoding center, the peptidyl-transfer center (PTC), and the inter-subunit bridges. Methylations and pseudouridylations are the most frequent of these rRNA modifications, which in bacteria are made by site-specific enzymes that act at different stages of ribosome assembly. Although high-resolution structures of ribosomes are available (1, 2), the precise contributions of rRNA modification to ribosome structure and function remain largely enigmatic.

In the *Escherichia coli* ribosomal large subunit (LSU), 13 out of the 36 known MNs(3) are located around the peptidyl transferase center in domain V of 23S rRNA, within 25 Å of the catalytically essential A2451 residue (Fig. 1). These include five pseudouridins (Ψ2457, Ψ2504, Ψ2580, Ψ2604, and Ψ2605), three 2’O ribose methylations (Gm2251, Cm2498, and Um2552), and three base methylations (m^7^G2069, m^2^G2445, and m^2^A2503), as well as one dihydrouridine (D2449) and one 5-hydroxycytidine (ho^5^C2501) (1). The genes encoding the corresponding enzymes are all known, but RdsA (D2449), which was only identified recently (4), is not included in this study. Recently, hypoxia-induced methylation at carbon 5′(*S*) of ribose moieties of dihydrouridine at position 2449 (D^5S^m2449) and 2′-*O*-metylcytidine at position 2498 (Cm^5S^m2498) by RlmX was identified (5). Ten MEs make 12 modifications in the domain V and one in the domain II (Ψ955) of 23S rRNA (Supplementary Table S1). Across species, highly conserved modifications around the PTC are Gm2251, Um2552, Ψ2457 and ho^5^C2501. Interestingly, there are also several examples of conserved tertiary interactions in ribosome structures, where stabilizing methylations occur on different nucleotide interaction partners across kingdoms of life (6).

**Figure 1.**
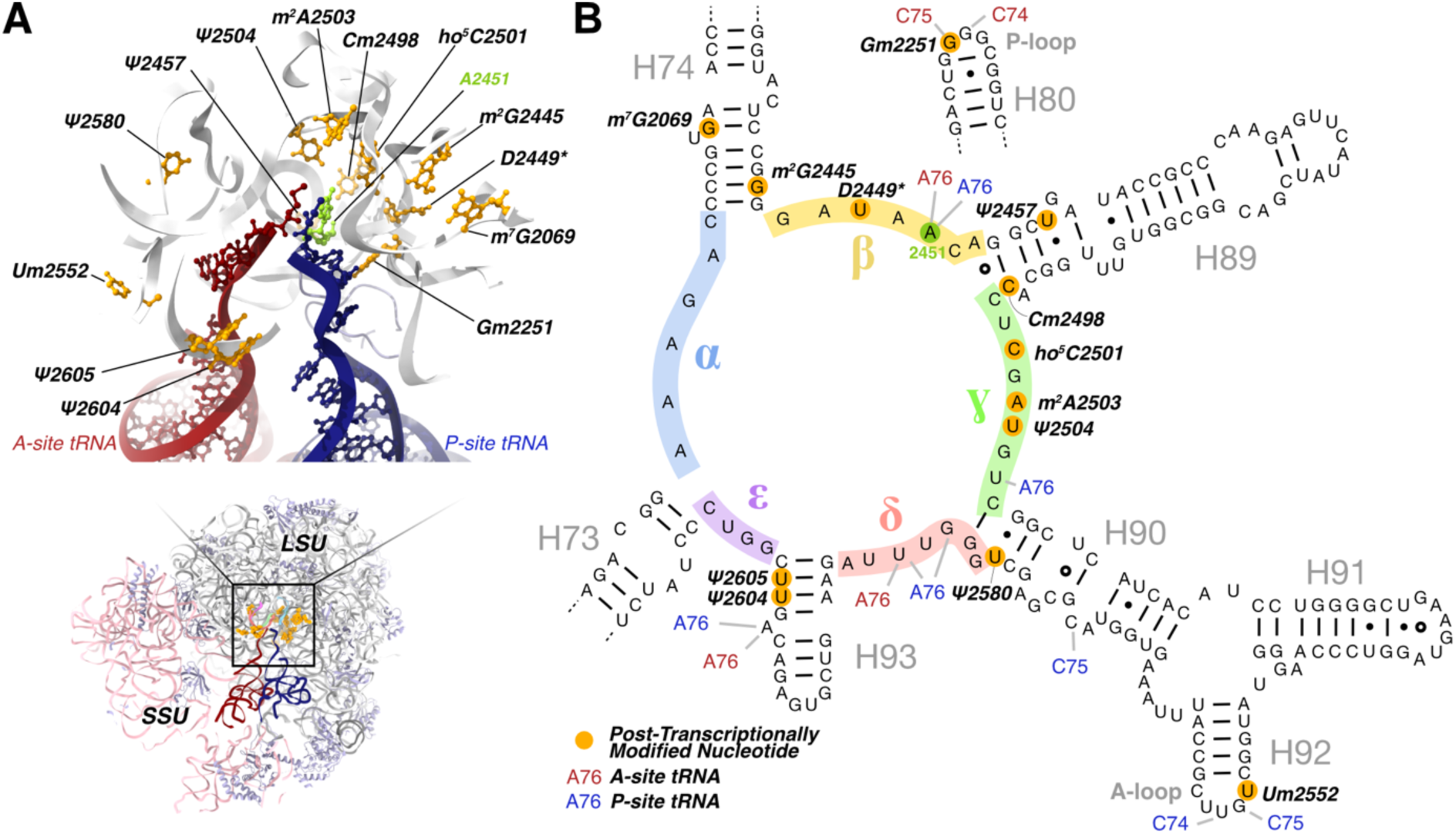
Post-transcriptional modifications at the PTC **A.** Modified nucleotides (orange) within 25 Å of the catalytic residue A2451 (green) to which none have direct contact nor to the tRNAs. **B.** Secondary-structure diagram of the PTC region of 23S rRNA. Sections of the PTC ring are labelled with Greek letters α–ε according to Supplementary Table S3. Nucleotides in contact with tRNA residues in PDB 8EMM are indicated. *D2449 is modified in all ribosomes in this study.

The functions of MNs have primarily been studied using KO strains. Single knock-outs of rRNA modification enzyme genes, with the exception of RlmE, which 2’O methylates U2552, do not significantly affect bacterial growth or translation (7, 8). In contrast, simultaneous depletion of nine or more enzymes leads to severe defects in growth and ribosome assembly (9). Interestingly, deletion of all seven rRNA-specific pseudouridine synthase genes results in only a weak phenotype in *E. coli* (*10*). However, deletion of ten MEs targeting domain V of 23S rRNA is possible, but the resulting strain shows severe growth retardation, defects in LSU assembly and cold sensitivity (9).

High-resolution structural studies have shown how MNs contribute to stabilizing the native rRNA structure in *e.g. E. coli* (1, 11) and human ribosomes (2). Generally, pseudouridines tend to form water-mediated hydrogen bonds with phosphate oxygens of the same nucleotide, thereby stabilizing the rRNA backbone (11), while base methylation increases base stacking propensity (12). Methylations are frequently found in regions of tertiary interactions.

Two conserved 2’O methylated nucleotides are located at the A (Um2552) and P (Gm2251) loops of 23S rRNA. Deletion of the RlmE gene, which specifically adds the Um2552 methylation results in decreased growth rate and major defects in LSU assembly (13–16). Although the rate of dipeptide formation between fMet-tRNA and Phe-tRNA is similar on the *ΔrlmE* and WT ribosomes, the rates of EF-G dependent translocation and ribosome subunit association are reduced in *ΔrlmE* ribosomes (16). The strong phenotype caused by RlmE depletion is further aggravated by deletion of the pseudouridine synthase RluC, which modifies positions 955, 2504, and 2580, especially at lower temperatures (17). In contrast, single deletion of *rlmB* (responsible for Gm2251) does not affect bacterial fitness or ribosome assembly (18).

The clustering of MNs around the PTC argues for their functional importance. Nevertheless, MNs are not strictly required for the peptidyl transferase (PTase) reaction, as *in vitro* transcribed unmodified 23S rRNA of thermophilic bacteria (19, 20) and *E. coli* (19) can catalyze peptide bond formation. However, the rate of the PTase reaction with unmodified 23S rRNA has not been analyzed. Despite detailed knowledge of the local interactions of MNs in the mature ribosome, the combined effects of MNs on rRNA folding and ribosome function remain poorly understood. Here, we analyze peptide bond formation on the ribosomes lacking chemical modifications in 23S rRNA around the PTC. Ribosomes lacking 11 or 12 MNs in the domain V of 23S rRNA catalyse peptide bond formation at a significantly reduced rate compared to the WT ribosomes and exhibit an increased entropic penalty of the reaction. Modified nucleotides confer thermotolerance to ribosomes, indicating stabilization of the native structure. Using single-particle cryo-EM analysis of programmed 70S ribosomes lacking 12 or 13 MNs, we show that MNs limit the structural heterogeneity of 23S rRNA, particularly its inherent propensity to form alternative stacked conformations. Consequently, ribosomes with a hypomodified PTC region display lower occupancy of A-site tRNA and of CCA ends correctly positioned for peptidyl transfer.

## MATERIAL AND METHODS

### Ribosome preparation

*E. coli* strains *Δ9*, *Δ10*, and *Δ9+RlmE* are described in (9). The *ΔrlmE* strain was constructed by P1 transduction using *ΔrlmE* of the Keio collection as a donor and *MG1655* as the recipient. Bacteria were cultivated in 2xYT (16 grams tryptone, 10 grams yeast extract, and 5 grams NaCl *per* 1 liter distilled water with pH adjusted to 7.0) or LB media supplemented with relevant antibiotics, and inoculated with a single colony and grown to mid-log phase. Cells were collected by low-speed centrifugation and resuspended in OV-10 lysis buffer (100 mM KCl, 50 mM Tris-HCl pH 8.0, 6 mM Mg(OAc)_2_, 16% sucrose, and 6 mM 2-mercaptoethanol) supplemented with DNase I (10 U/ml)). Cells were disrupted with glass beads using Bertin Precellys24 Tissue Homogenizer (3 cycles of 60 sec. on/off 6000 rpm at 4°C). Lysate was clarified by centrifugation (13000 rpm for 15 min at 4°C) and diluted 1:1 with OV-10 lysis dilution buffer (100 mM KCl, 20 mM Tris-HCl pH 8.0, 10 mM Mg(OAc)_2_, and 6 mM 2-mercaptoethanol). 80 (A_260_) units of lysate were layered onto a 10% to 30% (w/w) sucrose gradient in OV-10 lysis dilution buffer. Ultracentrifugation was carried out (ω^2^t=3.0 x10^11^ rad/sec) at 4°C using a Beckman Coulter SW-28 rotor. The ribosome profile was determined by continuous monitoring of absorbance at 260 nm. Fractions corresponding to 70S ribosomes and free 50S subunits were collected and stored at -80C°.

The 50S and 30S ribosomal subunits were isolated from 70S ribosomes under dissociating conditions. Briefly, 70S ribosome pellets were resuspended in OV-1 buffer (100 mM KCl, 20 mM Tris-HCl pH 8.0, 1 mM Mg(OAc)₂, and 6 mM 2-mercaptoethanol). A total of 80 A₂₆₀ units of 70S ribosomes were layered onto a 10–25% (w/w) sucrose gradient prepared in the same buffer and subjected to ultracentrifugation at 4 °C using a Beckman Coulter SW-28 rotor (ω²t = 2.8 × 10¹¹ rad²/s). Following centrifugation, the 50S and 30S fractions were collected, and the Mg(OAc)₂ concentration was adjusted to 10 mM to promote subunit stabilization. The subunits were then pelleted by ultracentrifugation using a Beckman Coulter Ti45 rotor (ω²t = 3.0 × 10¹¹ rad²/s). Finally, the ribosomal subunits were resuspended in polymix buffer (20 mM Hepes–KOH pH 7.6, 95 mM KCl, 5 mM NH₄Cl, 0.5 mM CaCl₂, 5 mM Mg(OAc)₂, 8 mM putrescine, 1 mM spermidine, and 1 mM dithioerythritol).

### Kinetic measurements

All experiments were performed in polymix buffer at 37 °C. The kinetics of dipeptide ([³⁵S]fMet-puromycin) and tripeptide (fMet[³H]Phe-puromycin) formation were analyzed using a RQF-3 rapid quench-flow apparatus (KinTek Corp., Austin, TX). The 30S initiation complex (30SIC) was assembled by incubating 30S ribosomal subunits (1 μM) with 2.5 μM A52 mRNA (CAA UUA AGG AGG UAU ACU AUG UUC GUA GCG AGC UAA; Microsynth), 2 μM [³⁵S]fMet-tRNAᶠᴹᵉᵗ (6000 dpm/pmol), and initiation factors IF1, IF2, and IF3 (each at a two-fold molar excess relative to ribosomes) at 37 °C for 15 minutes.

For the dipeptide assay, pre-formed 70S initiation complexes (70SIC) were assembled by incubating the 30SIC (1 μM) with 0.5 μM 50S subunits at 37 °C for 15 minutes. The complexes were isolated by centrifugation through a 1 ml sucrose cushion (20% w/w sucrose in buffer A) at 260,000 × g for 2 h using an Optima MAX-XP ultracentrifuge (Beckman Coulter). The resulting ribosome pellets were resuspended in polymix buffer, shock-frozen in liquid nitrogen, and stored at -80 °C.

The rate of [³⁵S]fMet-puromycin formation (dipeptide assay) was determined by rapidly mixing equal volumes of 1 μM 70SIC and preincubated 20 mM puromycin at 37 °C for defined time intervals. Reactions were quenched by the addition of 17% formic acid. To hydrolyze the tRNA, 150 μl of 6 M KOH was added, followed by neutralization with 500 μl of 1.5 M Tris-HCl (pH 8.8). The reaction products were extracted with 1 ml of ethyl acetate, and [³⁵S]fMet-puromycin was quantified from the organic phase by scintillation counting.

In the tripeptide assay, the rate of peptide bond formation between dipeptidyl-tRNA (fMet[^3^H]Phe-tRNA^Phe^) and puromycin was measured on post-translocation 70S initiation complexes (post70SIC). Briefly, 70SIC and elongation mixture (EM) were formed separately. 70SIC was formed by incubating 30SIC (1 μM) with purified 50S subunits (0,75 μM) at 37 °C for 15 min. EM contained 20 μM EF-Tu, 5 μM EF-G, 10 μM tRNA^Phe^, 1 μM [^3^H]phenylalanine ([^3^H]Phe specific activity 5000 dpm/pmol), and Phe-tRNA synthetase, supplemented with 1 mM ATP, 1 mM GTP, 10 mM phosphoenolpyruvate, and 0.05 mg/mL pyruvate kinase for energy regeneration. Post-translocation 70SIC complex carrying fMet[^3^H]Phe-tRNA^Phe^ in the P site was formed by mixing equal volumes of 70SIC and EM, incubated at 37℃ for 15 min and isolated by centrifugation through 1 ml sucrose cushions (20% sucrose w/w in buffer A) at 260,000g for 2 h (Optima MAX-XP, Ultracentrifuge; Beckman Coulter). Ribosome pellets were dissolved in polymix buffer, shock-frozen in liquid nitrogen, and stored at –80°C.

Formation of fMet[^3^H]Phe-Pmn was measured upon rapid mixing of equal volumes (15 μL) of purified pre-translocation 70SIC complex and Pmn (f.c. 20 mM) in the quench-flow apparatus. After the desired incubation time, the reaction was quenched with 17% formic acid, tRNA was hydrolyzed by the addition of 150 μl 6M KOH, neutralized by the addition of 500 μl 1 M Tris pH 8.0, and the tripeptide fMet[^3^H]Phe-Pmn was analyzed by extraction with ethyl acetate and subsequent scintillation counting. Note that unreacted [^3^H]Phe remains in the water phase at alkaline pH. The time courses were fitted to single-exponential kinetics according to the formula: F^fMet-Puro^ = F0*(1 − exp[−k_obs_ * t]), where F^fMet-Puro^ denotes the fraction of the fMet-puromycin formed, F0 is the maximal amount of the puromycin-reactive 70S complexes, and k_obs_ is the apparent first-order rate constant.

### Thermal inactivation assay

20 picomoles of 50S ribosomal subunits were incubated for 30 minutes in OV-10 buffer (100 mM KCl, 20 mM Tris-HCl pH 8.0, 10 mM Mg(OAc)₂, and 6 mM 2-mercaptoethanol) at various temperatures (37 °C, 42 °C, 49 °C, and 60 °C), followed by gradual cooling to 37 °C over a 30-minute period. Peptidyl transferase (PT) activity was assessed using the [³⁵S]fMet-puromycin assay by measuring the reaction product at the 2.5-minute time point. Briefly, 15 pmol of 30S subunits were preincubated with initiation factors, A52 mRNA, and [³⁵S]fMet-tRNAᶠᴹᵉᵗ for 20 minutes at 37 °C to form the 30S initiation complex. Subsequently, 10 pmol of the thermally treated 50S subunits were added. The reaction was initiated by the addition of puromycin (f.c. 10 mM), and the formation of [³⁵S]fMet-puromycin was measured after 2.5 minutes.

### Cryo-EM sample preparation

50S subunits were isolated from 70S ribosomes of *Δ9*, *Δ10,* and *ΔRlmE* strains as described above. 50S subunits (final concentration 1 µM) from deletion strains (Δ9, Δ10, and ΔRlmE) were assembled with wild-type 30S subunits (2 µM) into 70S ribosomes in the presence of uncharged tRNA^fMet^ (4 µM), tRNA^Phe^ (4 µM), and synthetic mRNA (8 µM) containing a Shine-Dalgarno sequence followed by an initiator codon and a phenylalanine codon (5’-caauuaaggagguauacuAUGUUCAAAGUACGAUCAAUCU ACGUAUAAUAAAAGAAAAGAAAAGAAAAGGACAUCACAGAUUAAcgcacaaaaaaaaaaaaaaaaaaa aa-3’). The assembly reactions were carried out in RS-6 Buffer (20 mM Tris-HCl, pH 7.5; 100 mM NH₄Cl; 6 mM magnesium acetate; 1 mM DTT) in a total volume of 150 µL at 37 °C for 20 minutes. 70S ribosomes were purified by 10–25% sucrose gradient centrifugation in RS-6 Buffer using an SW28 rotor (Beckman Coulter; *w²t* = 2.7 × 10¹¹). The 70S-containing fractions were pooled, and the magnesium acetate was adjusted to 10 mM, and sucrose was removed by dilution and concentration using a 100 kDa MWCO Amicon Ultra centrifugal filter unit (Millipore). The final ribosome storage buffer (RS Buffer) consisted of 2 mM Tris-HCl (pH 7.5), 100 mM NH₄Cl, 10 mM magnesium acetate, and 1 mM DTT. To ensure high tRNA occupancy, 1 µM of each uncharged tRNA was freshly added to the sample before freezing.

The samples were incubated for 30 seconds on QuantiFoil R 2/2 300 mesh copper grids coated with a continuous layer of 2-nm amorphous carbon before blotting for 4 seconds and plunge frozen in liquid ethane using a VitroBot mark IV (Thermo Fisher Scientific, Waltham, MA, USA) kept at 4 °C and 95% humidity. The grids were activated using a EasiGlow (TedPella Inc, Redding, CA, USA) set to 20 mA for 20 seconds in 0.4 mBar residual air.

### Cryo-EM data collection

Grids were screened on a Glacios (Thermo Fisher Scientific) microscope. The Δ9 and Δ10 data sets were collected on a Titan Krios G3 (Thermo Fisher Scientific) equipped with a K2 (Gatan, Pleasanton, CA, USA) camera at 105,000x magnification (equivalent to a calibrated pixel size of 0.8215 Å/pixel in the sample plane using) with a BioQuantum (Gatan) energy filter with a slit width of 20 eV and with a total dose of 30.0 e/Å^2^ saved as 20 frames. The ΔRlmE data set was collected on a Titan Krios G3 equipped with a K3 camera (Gatan) and BioQuantum energy filter at a nominal magnification of 105,000x (calibrated to 0.8228 Å/pixel) and an energy filter slit width of 20 eV and with a total dose of 40.6 e/Å^2^ saved as 39 frames.

### Cryo-EM data processing

Data was processed using Relion (21) and CryoSPARC (22) according to the conventional single-particle processing pipeline. In short, the movies were motion corrected using the Relion 3 (23) implementation of the MotionCor2 algorithm (24) with 5×5 patches. The initial CTF parameters were estimated using Gctf (25), and particles were picked by matching round (Laplacian of Gaussian) templates to features in the micrographs. Particles were extracted, binned 4×4, and several rounds of 2D classification were used to remove non-ribosomal particles. A consensus 3D reconstruction of the 70S ribosome was obtained after iterative rounds of 3D reconstruction and estimation of optical parameters using the Relion 4 algorithm for per-particle defocus and higher order aberrations (21), and particle polishing (23). Focused classification without alignment into 6 classes within a soft-edged mask over the tRNA binding sites, based on the consensus 70S reconstruction after subtracting the signal of the LSU and the SSU segregated particles into classes with tRNA density in the A, P, and/or E sites. Relevant APE, AP, PE & E classes were reconstructed. The occupancy of tRNA in each class was estimated based on the local loss of contrast in the reconstructions using Occupy (26). The polished particles from the consensus reconstruction were imported into CryoSPARC 4 and were 3D classified into 10 classes without alignment based on a 40-Å spherical soft-edged mask over the PTC region after subtracting the signal of the LSU and the SSU. Maps were globally sharpened using a negative B-factor based on fitting to the Guinier plot and low-pass filtered based on the estimated FSC resolution. The ΔRlmE dataset was processed in the same way, but entirely in CryoSPARC 4, using the Patch Motion for motion correction, the Patch CTF algorithm for initial CTF estimation, and the blob picker for particle picking. The magnification for each dataset was calibrated by rigid-body docking PDB ID 7K00 (1) in the consensus maps in ChimeraX (27) and maximizing the map-model correlation by changing the voxel size.

### Model building

Atomic structures were generated based on PDB IDs 8CF1, 8CGJ, and 8CGK (28) and 7K00 (1). The chains were rigid-body docked into the consensus ΔRlmE structure using ChimeraX and Coot (29) version 0.9.8.95. Residues outside of density or with validation issues were interactively refined or rebuilt using Coot. The tRNA^Phe^ in the A-site, uL5 and 23S helices H25, H42-45 and H74-76 were was modeled based on an AlphaFold 3 (30) predictions. The E-site tRNA was modeled as tRNA^fMet^, although it is occupied by a mixture of tRNA^fMet^ and tRNA^Phe^. Modelling the consensus Δ9 and Δ10 structures were based on the refined consensus ΔRlmE model. Atomic models for residues close to the PTC-region (45-Å distance from A2451) were modeled in the maps for ΔRlmE, Δ9 and Δ10 classes from the focused classification of the PTC-region. The final structures were refined using Servalcat (31) version 0.4.77 with jelly-body restraints. The CCP4 monomer library (32) was used to restrain bond lengths, angles, torsions, planarity, and chirality for standard residues, while restraints for modified residues were generated by Grade2 using the Grade Web Server (https://grade.globalphasing.org, Global Phasing Ltd, Cambridge, UK). The models were validated using Coot, Phenix (33) version 1.21, and MolProbity (34) as distributed in the Phenix package (33). Refinement and validation information is summarized in Supplementary Table S5.

### Statistical analyses

Statistical analysis of experimental data was done using the ordinary two-way ANOVA tests and multiple comparison was done using uncorrected Fisher’s LSD tests.

## RESULTS

### Pre-Steady-State Kinetics of Peptide Bond Synthesis on Ribosomes

We set out to analyse the importance of 23S rRNA modifications during peptide bond formation on the ribosome. To this end, ribosomes were isolated from *E. coli* strains lacking either 11 (*Δ9*) or 12 (*Δ10*) modifications around the PTC, which show accumulation of incompletely assembled ribosomes (9). Since the aim was to analyse the PTase activity that supports viability of the strains, we isolated fully assembled 70S ribosomes, performed subunit dissociation in low Mg^2+^ to isolate the hypomodified 50S subunits, and combined these with WT 30S subunits in all experiments. To determine the kinetic parameters of peptide bond synthesis, we employed an assay system utilizing puromycin (Pmn) as acceptor substrate and peptidyl-tRNA analogs (fMet-tRNA in the dipeptide assay and fMetPhe-tRNA in the tripeptide assay) as the donor substrate. We selected the puromycin reaction as a model system for peptide bond synthesis, as this system does not involve the multistep accommodation of aa-tRNA to the ribosomal A site (35). In this assay, 70S complexes (70S*fMet-tRNA*mRNA or 70S*fMetPhe-tRNA*mRNA) were formed in the presence of purified translation initiation and elongation factors, purified via sedimentation through a sucrose cushion and kept at -80 °C until use. The same assay system was previously used to define the effect of mutations in both 23S rRNA and ribosomal protein L27 on the rate of peptide bond formation (36, 37).

In the dipeptide assay, the rate constant k_obs_ for the PTase reaction on WT ribosomes was 1.39 ± 0.13 sek^-1^ (Fig. 2A, B) at 37 °C, in agreement with earlier measurements (37, 38). Ribosomes of *Δ9* and *Δ10* strains catalyzed dipeptide formation with rates of 0.62±0.14 and 0.42±0.13 sek^-1^, respectively (Fig. 2A, B). 50S subunits of the *Δ10* strain expressing plasmid-encoded RlmE (*Δ10+rlmE*), containing the same 23S rRNA modification pattern as Δ9 ribosomes, exhibited k_obs_ of 0.57±0.09 (Fig. 2A, B). Thus, hypo-modified ribosomes catalyze peptide bond synthesis 2.2 – 3.3 times slower than fully modified WT ribosomes, and RlmE appears to stimulate the PTase reaction slightly. We conclude that some of the 23S rRNA modifications around the PTC stimulate the rate of the PTase reaction.

**Figure 2.**
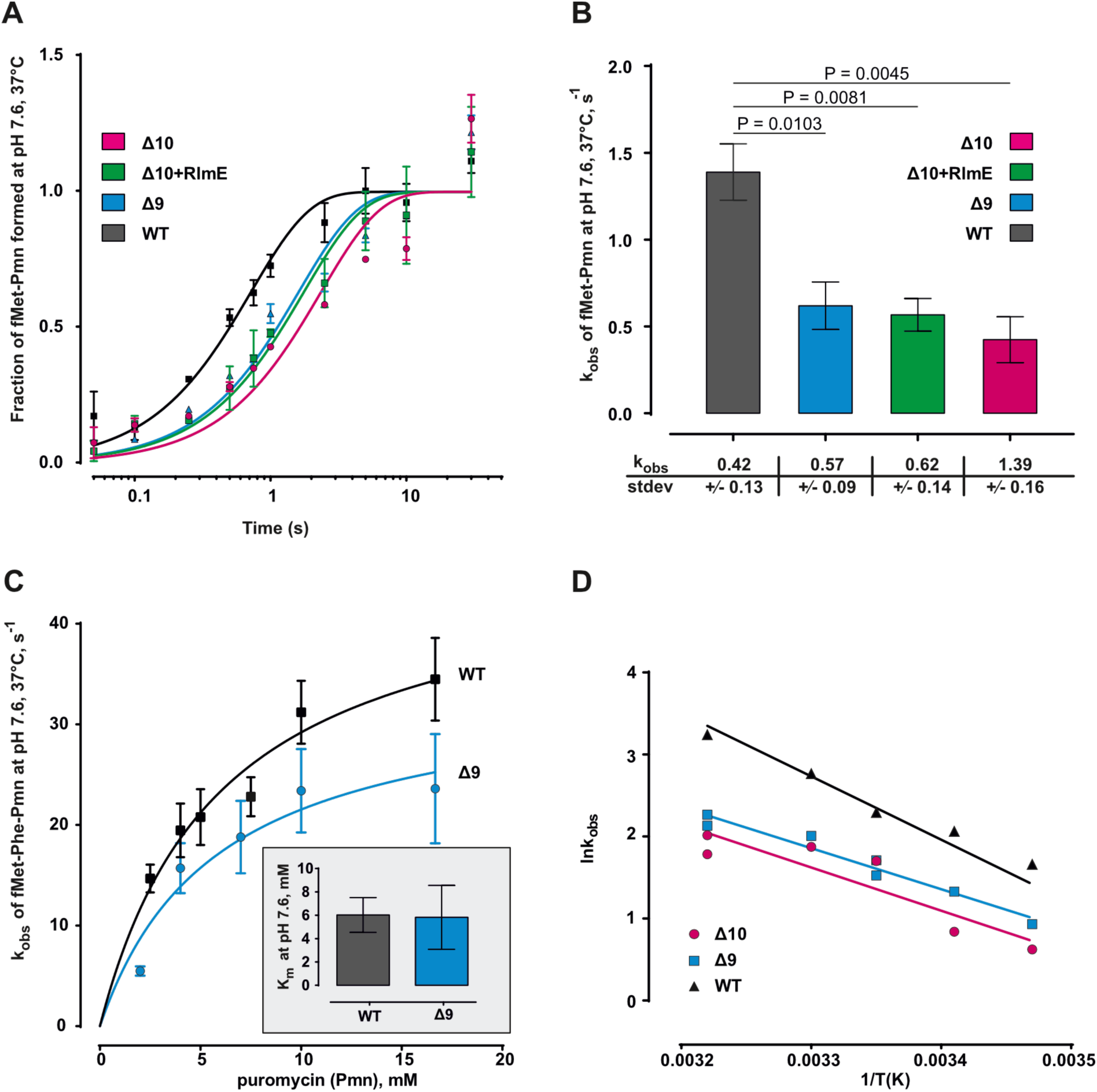
Modifications in the domain V of 23S rRNA stimulate peptide bond formation in the fMet-tRNA and puromycin reaction (Dipeptide assay) and in the fMetPhe-tRNA and Pmn reaction (Tripeptide assay). Ribosomal 50S subunits were isolated from 70S ribosomes of WT, Δ9, Δ10, and Δ10+RlmE strains. In dipeptide assay fMet-tRNA*mRNA*70S ribosome complex (70 SIC) was mixed with an equal volume of 20 mM Pmn in a quench-flow machine and incubated for 0.1 – 10 seconds. In tripeptide assay of fMetPhe-tRNA*mRNA*70S complex was reacted with Pmn. **A.** Time course of dipeptide PTase reaction. **B.** k_obs_ values of PTase reaction with various ribosomes (dipeptide assay). **C.** Titration of fMetPhe-tRNA*mRNA*70S complex with Pmn at pH 7.2. Inlet K_M_ values of the PTase reaction on the ribosomes of WT and Δ9 strain. **D.** Effect of temperature (15 – 37 °C) on the PTase reaction using tripeptide assay shown as Arrhenius plot.

The PTase reaction rates between fMet-tRNA and Pmn on WT, Δ9, and Δ10 ribosomes were measured at pH 7.2 and 7.6 (Supplementary Fig. S1). The PTase reaction was about 3 times faster on WT ribosomes than on Δ10 ribosomes at both pH values. Ribosomes of the Δ9 strain exhibited slightly higher activity than the Δ10 ribosomes (Supplementary Fig. S1). The reaction rate constant (k_obs_) increased 3.7-fold at the higher pH for all three ribosome variants (Supplementary Fig. S1), consistent with previously published data with puromycin as acceptor substrate (35). Thus, the lack of modifications in domain V of 23S rRNA does not appear to affect the pH dependence of the PTase reaction.

Peptide bond synthesis between fMet-tRNA and Pmn is slow as compared to the translation elongation rate *in vivo* (1.4 vs 20 s^-1^) (Fig. 2B) (35). However, with dipeptidyl-tRNA (fMetPhe-tRNA) as donor substrate, the PTase reaction rate is more than an order of magnitude higher (35, 36, 39). Therefore, the tripeptide assay was used to determine the effect of 23S rRNA modification on the rate of PTase reaction. Indeed, the maximal rate of peptidyl-Pmn synthesis was about 35 s^-1^ on the WT ribosomes at pH 7.6 and saturating Pmn concentration (Fig. 2C). In agreement with the earlier observations (35, 36, 39), this is about 25 times faster than with fMet-tRNA as a donor substrate, likely explained by additional hydrogen bonds between dipeptidyl-tRNA and 23S rRNA that stabilize the active PTC conformation (40). To test whether the puromycin affinity to the ribosomal A-site is affected by the absence of modifications, the tripeptide assay was performed at various Pmn concentrations on WT and Δ9 ribosomes (Fig. 2C). For this experiment, ribosomes from the Δ9 strain were used, as in this strain, ribosome assembly is not significantly affected at 37 °C (9), but as described above, ribosomes from Δ9 and Δ10 showed similar rates in the dipeptide assay. For WT and Δ9 ribosomes, the K_M_ value was 6 mM for both WT and Δ9 ribosomes (Fig. 2C). Thus, the PTC modifications do not affect the affinity of puromycin to the ribosomal A site.

Temperature dependence (15 – 37 °C) of peptide bond formation on the ribosomes from the WT, Δ9, and Δ10 strains was analysed using the tripeptide assay. This assay was done at pH 7.2 to improve the solubility of puromycin at low temperatures. Both Δ9 and Δ10 ribosomes catalyse the PTase reaction at a similar rate, slower than the rate on WT ribosomes at all temperatures (Supplementary Table S2). However, the difference in k_obs_ values between the WT and hypo-modified ribosomes increases with increasing temperature. Thus, the functional significance of the 23S rRNA modifications in peptide bond formation appears stronger at higher temperatures. At 37 °C, the WT ribosomes catalyse peptide bond formation about three times faster than hypo-modified ribosomes, similar to the results of the dipeptide assay. These results suggest that 23S rRNA modifications around the PTC are important for stabilizing the binding of either substrate or a reaction intermediate to the ribosome.

The first order rate constants of the tripeptide assay were used to calculate the thermodynamic parameters of the PTase reaction based on the Arrhenius plot (Fig. 2D; Table 1). The values of parameters ΔH and TΔS of the reaction of WT ribosomes were very similar to the previously observed values (41). Comparison of WT and hypo-modified ribosomes demonstrates that the ribosomes of *Δ9* and *Δ10* strains exhibit similar values for all thermodynamic parameters but deviate significantly from WT ribosomes. Surprisingly, the activation energy (Ea) and enthalpy (ΔH) are higher on the WT ribosomes than on the Δ9 and Δ10 ribosomes, suggesting that modifications around the PTC increase the activation energy barrier of the reaction. On the other hand, the TΔS value of the PTase reaction is significantly higher on the Δ9 and Δ10 ribosomes. Thus, the modifications around the PTC reduce the entropic penalty of peptide bond formation on the ribosome. Reduced enthalpy and increased entropy can result from a weakened binding of substrates to the PTC. This idea is compatible with the hypothesis that one or more rRNA modifications near the PTC are crucial for the orientation of substrates during peptidyl transfer. Alternatively, rRNA modifications can stabilize the binding of the reaction intermediate to the PTC.

**Table 1.** Thermodynamic parameters of the fMetPhe-tRNA reaction with Pmn (Tripeptide assay) on the ribosomes of *WT* and hypo-modified strains *Δ9* and *D10*.

|  | **E<sub>a</sub>** | **SD** | **ΔH** | **SD** | **ΔG** | **SD** | **TΔS** | **SD** |
| --- | --- | --- | --- | --- | --- | --- | --- | --- |
| **WT** | 15.53 | ±1.72 | 14.94 | ±0.62 | 16.09 | ±0.08 | 1.16 | ±0.69 |
| **Δ9** | 10.94 | ±1.01 | 10.35 | ±1.01 | 16.44 | ±0.05 | 6.10 | ±1.05 |
| **Δ10** | 10.59 | ±2.43 | 10.00 | ±2.43 | 16.45 | ±0.1 | 6.45 | ±2.53 |

### Modifications stabilize the 50S subunit structure

The lacking rRNA modifications around the PTC appears to affect the rate of peptide bond formation more at higher temperatures (Supplementary Table S2; Figs. 2D and 3), suggesting a possible problem with ribosome stability. If rRNA modifications stabilize the ribosome structure, a lack of modifications is expected to cause reduced thermotolerance. To test this, the 50S subunits were first pre-incubated for 30 min at 37 °C, 42 °C, 49 °C, or 60 °C, followed by measurement of PTase activity. Pre-incubation of the WT 50S subunits at any temperature did not affect their ability to catalyse peptide bond formation (Fig. 3), in agreement with (42). In contrast, pre-incubation of the Δ10 50S at 49 °C reduced PTase activity by 10%, and at 60 °C by 60% (Fig. 3). The 50S subunits of Δ9 and Δ10+RlmE strains exhibited similar stability at elevated temperatures as Δ10, albeit the reduction of PTase activity was slightly lower as compared to the Δ10 ribosomes (Fig. 3). These results demonstrate that the modifications around the PTC are important for LSU stability. Although RlmE is important for 50S assembly (13–16), its resulting methylation, Um2552, has little impact on the LSU stability, as evidenced by similar thermotolerance of the 50S subunits of Δ9 and Δ10 strains. The results demonstrate that the 23S rRNA modifications around the PTC protect ribosomes from heat inactivation and are important for ribosome structural stability. However, since heat inactivation of hypo-modified 50S subunits occurs during prolonged incubation at 49 °C or higher temperatures but not at 37 °C, it does not explain the reduced PTase activity of the Δ9 and Δ10 ribosomes at 30 °C and 37 °C.

**Figure 3.**
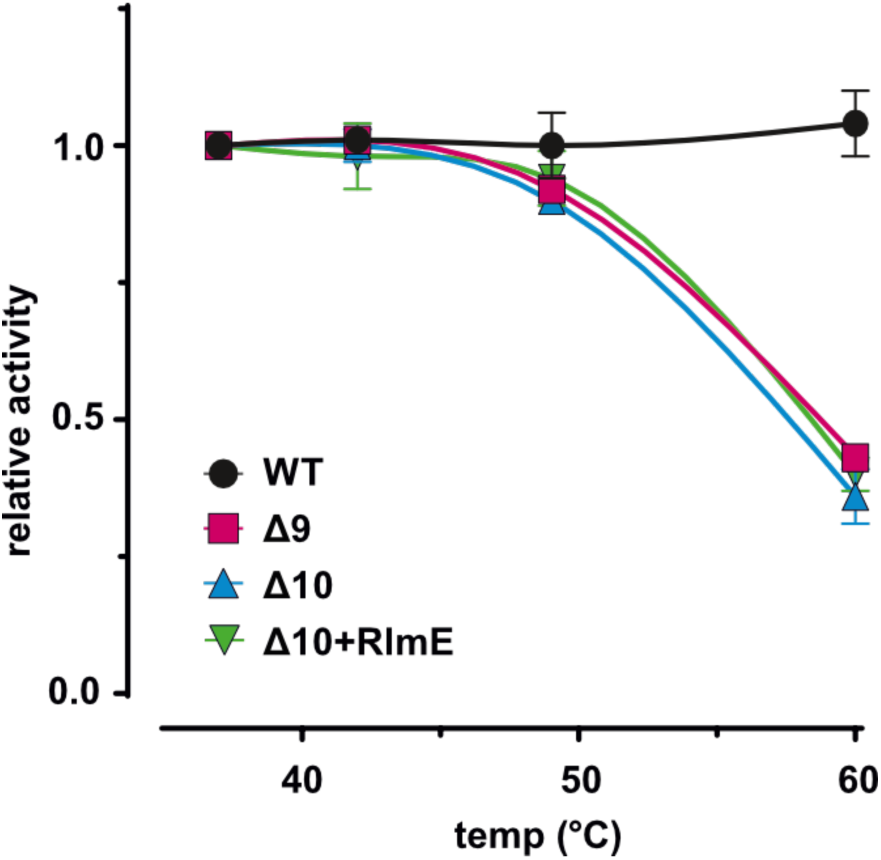
Effect of 23S rRNA modifications on the stability of 50S subunits. Ribosomal 50S subunits were preincubated at various temperatures, and 70SIC was formed with fMet-tRNA, and the active fraction of ribosomes according to PTase activity was determined using Pmn as an acceptor substrate.

### Structure determination of hypomodified ribosomes by cryo-EM

To shed light on the structural consequences of depleting the 70S ribosome of its native PTC-region modifications, we employed single-particle cryo-EM analysis. Grids were prepared with Δ9, Δ10, and, as a control, ΔRlmE 70S ribosomes programmed with a short synthetic mRNA, tRNA^fMet^ at the P-site, and tRNA^Phe^ at the A-site. Cryo-EM analysis resulted in consensus 70S ribosome reconstructions at better than 1.9 Å for all three samples (Supplementary Fig. S2–S4).

The structure of the ΔRlmE 70S ribosome is overall very similar to the WT *E. coli* 70S ribosome (1, 28). In contrast, the Δ9 70S structure, where the RlmE modification Um2552 is present has regions of weak density at the peptidyl transfer centre, resulting from flexibility and partial misfolding of 23S rRNA as well as r-protein uL4. Alternative 23S rRNA conformations are mainly observed in the PTC ring (regions α-ε, as defined in Fig. 1B and Supplementary Table S3). In particular, nucleotides 2058–2062 (in PTCα) and 2501–2506 (in PTCγ) show significant disorder with evidence of multiple conformations. The Δ10 70S ribosome shows additional disorder and misfolding, including also nucleotides 2581–2585 in PTCδ and the A-loop in H92. The CCA loop of the A-site tRNA also show several misfolded structures in all three samples (Fig. 4B–E and Supplementary Fig. S5).

**Figure 4.**
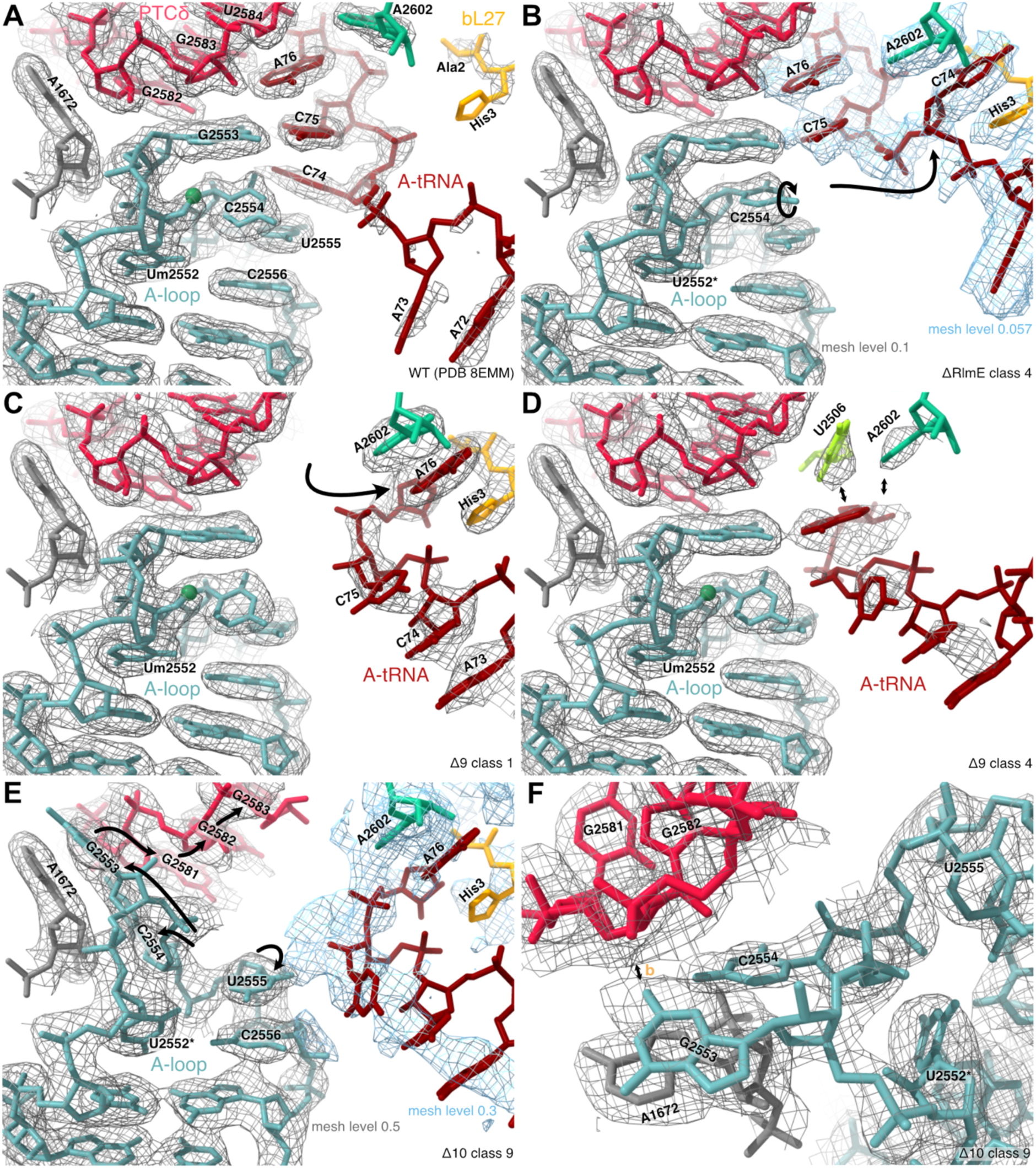
A-loop conformation with A-site tRNA. Segmented maps shown as mesh within 2 Å of residues. Supplementary Fig. S5 shows all PTC classes. **A.** Wild-type conformation (PDB 8EMM). The O2’ methyl group of Um2552 is indicated with a green sphere. **B.** ΔRlmE class 4 with C2554 rotated. C74 of the A-site tRNA (dark red) is sequestered between A2602 (mint green) and His3 of bL27 (orange). tRNA density shown at lower threshold as blue mesh. *U2552 unmodified. **C.** Δ9 class 1 with WT conformation of the A-loop. A76 sequestered between A2602 and His3 of bL27. **D.** Δ9 class 4 with WT conformation of the A-loop. U2506 (lime green) and A2602 (mint green) base-stack and block the binding-site of A76 (tRNA modelled similar to Δ9 class 1). **E.** Δ10 class 9 with flipped-out G2553 and U2554 stacking on A1672, which repositions 2581-2585 in PTCδ (cherry red). The CCA end is deflected (tRNA modelled similar to Δ9 class 1). **F.** Close-up of G2553 and U2554 in E from a different angle. The flipped-out bases clash with the WT conformation of PTCδ (non-native contact b, Supplementary Fig. S8).

The presence or absence of methylations could be confirmed directly in the maps at all nucleotide positions that were ordered in the consensus reconstructions (Supplementary Fig. S6). As a consequence of lacking methylation, several unmodified amine or hydroxyl groups are observed to coordinate structured water molecules (G2069, C2498 and U2552) or form non-canonical polar contacts (G2251 with D2449, discussed below). The non-planar base geometry of D2449 is resolved in ΔRlmE and Δ9 maps and the posttranscriptionally added hydroxyl group of ho^5^C2501 is clearly visible in ΔRlmE (Supplementary Fig. S6). The two isomers pseudouridine and uridine are isosteric and thus cannot be distinguished by shape in the maps, but we observe increased distance from C5 of uridine to the closest structured water in U2457, U2580 and U2605 (Supplementary Fig. S6).

Focused 3D classification on the tRNA binding pockets of the three reconstructions segregated the particles into classes depending on their tRNA occupancy in the A, P and E sites (Supplementary Table S4). Particles with A-site tRNA is fewer in Δ9 ribosomes compared to ΔRlmE and only 26% in Δ10. Even after classification, the tRNAs showed weaker density than the ribosome. Thus, tRNA occupancy was estimated based on local loss in contrast to be 45%, 35% and 21% in the A site, and 70%, 61% and 45% in the P site of the ΔRlmE, Δ9, and Δ10 ribosomes, respectively (Supplementary Fig. S7).

Diffuse density around the PTC indicated the presence of several conformations, and motivated further focused classification. Using a spherical mask around the PTC, it was possible to resolve multiple alternative conformations, in particular for Δ9. Notably, PTC classes with WT-like conformation also have high occupancy of A-site tRNA.

### Misfolding of the PTC ring

In the native WT structure, the three MNs of PTCγ form an intricate network of tertiary contacts with PTCα, PTCβ and PTCδ. In absence of these modifications, several of the modelled conformations show complete refolding of PTCγ, with disrupted and alternative tertiary contacts (Supplementary Fig. S8A). One such example is m^2^A2503 that in the native structure is intercalated in the PTCα stack while G2502 stacks with G2060 (Fig. 5A). These interactions are missing in several structures of Δ9 and Δ10 (Fig. 5B–D, Supplementary Fig. S9), resulting in misplacement of residues in the beta, gamma and delta loops that are important for positioning the tRNA substrates.

**Figure 5.**
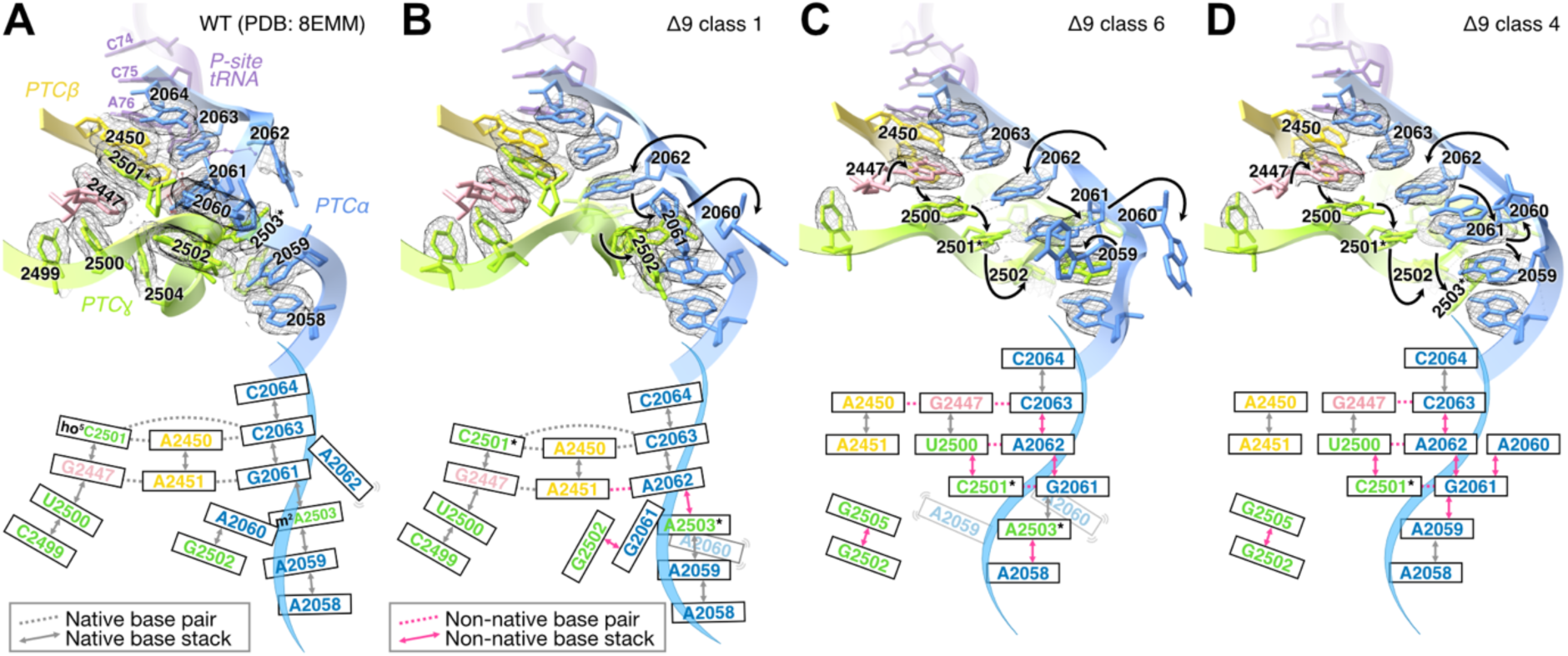
Misfolding of the PTC ring in 70S Δ9 ribosomes with A-site and P-site tRNA. Residues colored as in Fig. 1B. Segmented maps shown as mesh within 2 Å of bases. Supplementary Fig. S9 shows all PTC classes. Insets show schematic diagrams with bp or base triplets as dotted lines and stackings as double-headed arrows, with non-native contacts in magenta. **A**. WT conformation (PDB 8EMM). **B.** Misfolded Δ9 PTC class 1. A2062 displaces G2061. A2060 flips out and disturbs uL4 (Supplementary Fig. S10). **C.** Misfolded Δ9 PTC class 6. G2447 replaces C2501 (*unmodified, Fig. 6). **D.** PTCα forms an independent stack with only bp contacts to PTCγ and PTCβ.

In several conformations, the misfolded PTCα clashes with the 55-73 loop of r-protein uL4, inducing disorder with weak density for an alternative conformation (Supplementary Fig. S10). Importantly, this loop forms a restriction point in the exit tunnel, and its deletion has been shown to induce cold sensitivity, lead to the accumulation of assembly intermediates, and alter the sensitivity pattern to exit-tunnel binding antibiotics (43).

Another example is ho^5^C2501, which in the WT structure stacks between G2446 and G2447 and forms a base triplet with A2450 and C2063 (Fig. 6A). In the hypo-modified ribosomes, C2501 is destabilized by the loss of two polar contacts to D2449 and G2446, allowing it to reposition to base pair with G2061 (Fig. 5C, D). This reconfiguration also disrupts the neighbouring base-triplet between G2061, G2447 and the catalytically essential A2451.

**Figure 6.**
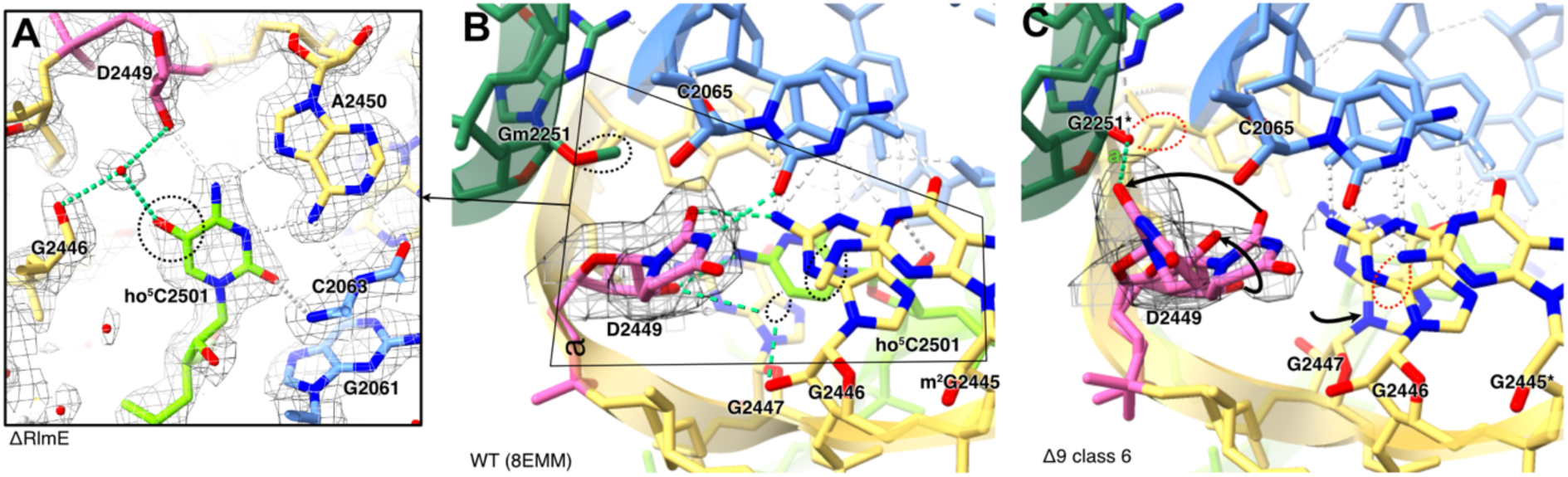
Confirmation of ho^5^C2501 and D2449. **A.** ho^5^C2501 (lime green) is in WT conformation in the consensus reconstruction of ΔRlmE. The posttranscriptionally added hydroxyl group (dashed ring) forms two water-bridged polar contacts (mint green dashed lines) to nearby ribose groups of G2446 and D2449. Additionally, ho^5^C2501 forms a base-triplet with A2450 and C2063. **B.** Wild-type conformation of D2449 (pink) with polar contacts (mint green dashed lines) to ho^5^C2501 (lime green), C2065 (light blue) and G2446 (yellow). Posttranscriptional modifications are circled with black dotted lines (ho^5^ of C2501 is not included in the model). **C.** Alternate conformation of D2449 flipped around in Δ9 class 6, stabilized by a hydrogen bond (mint green dashed line, non-native contact a, Supplementary Fig. S8) to the unmethylated O2’ oxygen of G2251 (dark green, *unmodified). C2501 is replaced by G2447 (yellow, *unmodified). Missing posttranscriptional modifications are circled with red dotted lines. Segmented map shown within 2 Å of D2449 in B and C.

In the absence of modifications in neighbouring nucleotides, the base of D2449, the only remaining PTC MN in Δ10, is shifted in several structures. In the most prevalent alternative conformation, the base of D2449 has shifted from anti to syn conformation, allowing it to hydrogen bond to the non-modified ribose hydroxyl group of G2251 (Fig. 6C and Supplementary Fig. S11). In WT ribosomes, this contact is prevented by the methylation of Gm2251 (Fig. 6B).

### Misfolding of the A loop affects binding of A-site tRNA

In the Δ10 ribosomes, the missing 2’O methylation of U2552 leads to misfolding of the A-loop (Fig. 4E, F). In the predominant conformation, G2553 and U2554 are flipped out and form a cis Hoogsteen-sugar basepair stacked on A1672. This conformational change explains the lower occupancy of A-site tRNA in D10, since G2553 and U2554 cannot form their normal hydrogen bonds to C74 and C75 of the A-site tRNA (Supplementary Fig. S5). The flipped-out bases clash with PTCδ, which is refolded such that G2581 stacks on G2582. As a consequence, many polar contacts between the A-loop and PTCδ are disrupted and PTCδ is largely disordered in most Δ10 classes, affecting *e.g.* G2583 and its contact with A76 of the A-site tRNA. A similar conformational change of G2553 and U2554 was previously observed in ΔRlmE 50S ribosomes (16) (Supplementary Fig. S5).

Notably, programmed ΔRlmE ribosomes show only minor differences from the WT conformation. In some classes, the base of U2554 is rotated and fully stacked between the neighbouring bases, clashing with C74 of the A-site tRNA, which is shifted to stack between His3 of bL27 and A2602 (Fig. 4B).

### Alternative stacking of the PTC perturbs the binding site of the A- and P-site tRNA acceptor ends

In the WT ribosome, U2506, U2585, and A2602 are dynamic in the absence of tRNA, but during translation adopt distinct conformations to position the aminoacyl and peptide groups of the tRNAs for peptidyl transfer (Fig. 7 and Supplementary Fig. S12). When PTCγ is perturbed, these nucleotides adopt several alternative conformations (Fig. 7B, C). In some classes, G2505 intercalates between U2506 and U2585 (Fig. 7B). In other classes, U2585 flips around to stack on U2584 while U2506 instead forms a stacking interaction with A2602, forming a barrier between the A76 nucleotides of the A- and P-site tRNAs (Fig. 7C). These misfolded conformations are incompatible with peptidyl transfer.

**Figure 7.**
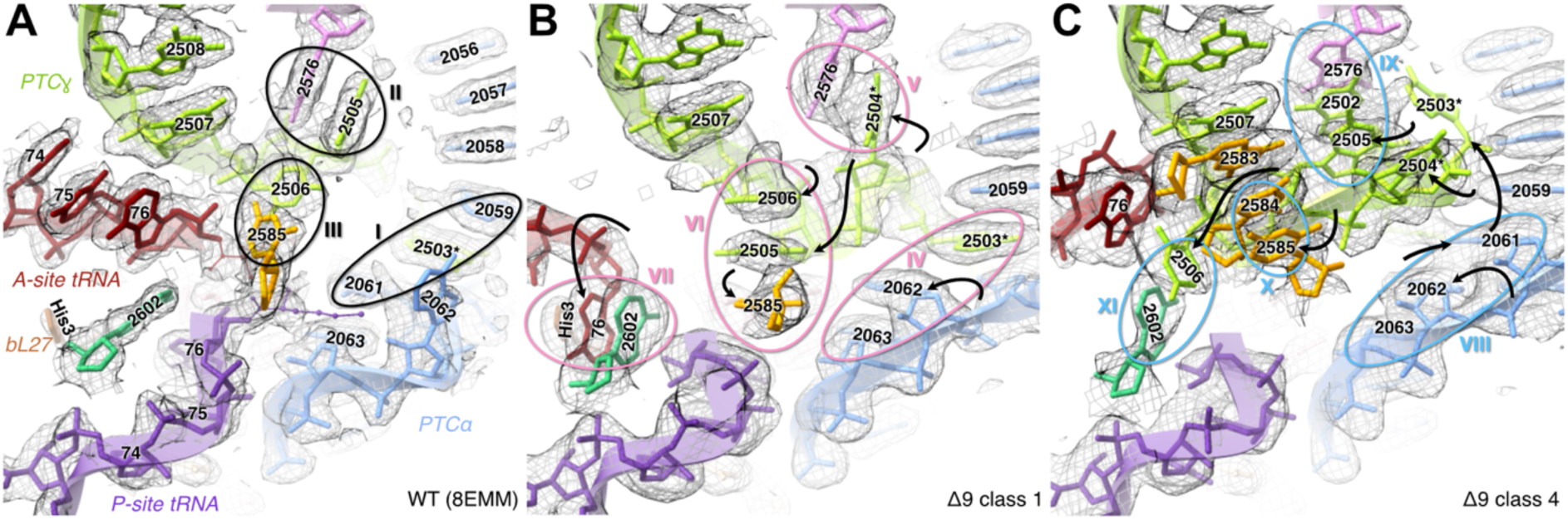
Alternative stacking at the PTC. Roman numbers refer to Supplementary Fig. S8B. Residues colored as in Fig. 1B. Segmented maps shown as mesh within 2 Å of residues in any of the classes. Supplementary Fig. S12 shows all PTC classes. **A.** WT conformation with bound tRNA (PDB 8EMM). PTCγ forms tertiary stacking interactions with PTCα (I), G2576 (H90 bulge, II), and U2585 (PTCδ, III). **B.** Misfolded Δ9 PTC class 1 with A2062 flipped in (IV, Fig. 7B), U2504 (*unmodified) replacing G2505 (V), G2505 intercalating between U2585 and U2506 (VI). A2602 stacks between A76 in the A-site tRNA and His3 in bL27 (VII, Fig. 4C). **C.** Misfolded Δ9 PTC class 4 with a novel stack G2576, G2502, and G2505 (VIII, Fig. 7D), U2585 stacking on U2584 (IX), and U2506 stacking with A2602, obstructing access between the A-site and the P-site (X, Fig. 4D).

### Modifications modulate the conformation of H89

H89, between PTCβ and PTCγ, is located close to the catalytic A2451 and the acceptor end of the A-site tRNA (Fig. 1). In Δ9 classes with no or weak density for A-site tRNA, we observe compression of one side of H89, near the modification sites of U2457 (WT psi) and C2498 (WT Cm), by up to 3 Å (Fig. 8C, D). The shift correlates with a conformational switching of the unpaired U2491, also seen in Δ10 (Supplementary Fig. S13). This switch has not been previously described, but mutations of U2491 were shown to affect PT activity (44). A similar but smaller deformation of H89, by about 1 Å, can be seen in WT ribosomes upon binding of A-site tRNA (Fig. 8A, B). This allows the phosphate of G2494, which base pairs with Ψ2457, to form a polar contact to His3 of bL27, and a magnesium-solvent bridge to catalytically essential A2451. This suggests that the MNs in H89 modulate the helix flexibility to prevent large deformation upon U2491-induced switching, assuring the correct geometry of the PTC.

**Figure 8.**
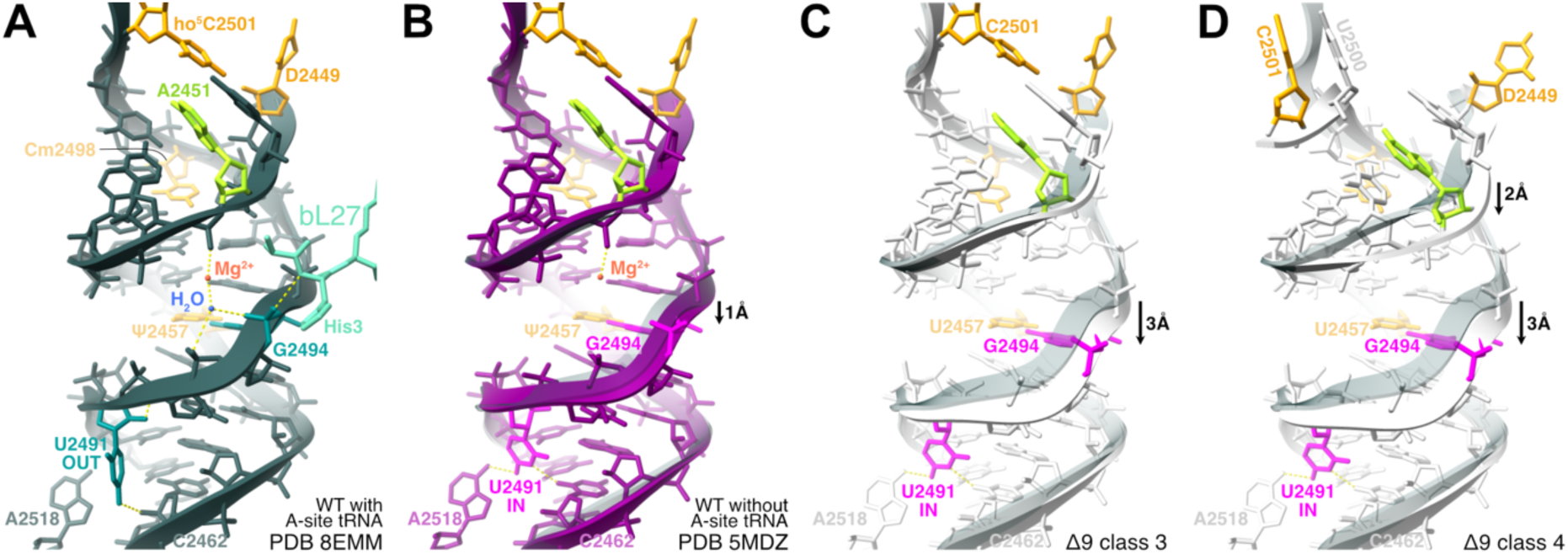
Switching of U2491 and deformation of H89. **A.** In WT ribosomes with A-site tRNA (PDB 8EMM), U2491 (teal) adopts an outward conformation (OUT) and hydrogen bonds (yellow dashed lines) to the ribose of C2462. G2494 (teal) has a solvent-bridged contact to A2451 (lime green) bridged by solvent and a hydrogen bond stabilizes the N-terminus of bL27 (mint green). Modified nucleotides show in orange. **B.** In WT ribosomes without A-site tRNA (PDB 5MDZ), U2491 (magenta) is tucked in and hydrogen bonds to the base of C2462 and to A2518 (IN). The helix is shifted by approximately 1 Å at G2494 relative to PDB 8EMM (semi-transparent dark green cartoon). **C.** Δ9 class 3 without A-site tRNA. G2494 is shifted by 3 Å relative to PDB 8EMM. **D.** Δ9 class 4 without A-site tRNA and misfolded PTC-ring. In addition to the shift of G2494 similar to class 4, catalytic nucleotide A2451 is displaced by approximately 2 Å relative to WT.

## DISCUSSION

Bacteria harbor genes for many enzymes dedicated to specific posttranscriptional rRNA modification, of which a large fraction cluster around the PTC. Here, we set out to combine kinetic and structural characterization of the Δ9 and Δ10 70S ribosomes, aiming to explain the collective effect of deletion of these rRNA modifications on the structure and catalytic activity and to thereby elucidate their combined function in WT ribosomes.

We were able to resolve an ensemble of misfolded PTC structures. This was technically possible since the main part of the ribosomal particles were well-ordered, guiding the particle alignment to high-resolution consensus reconstructions. The structural heterogeneity was limited to a well-defined region close to the PTC at the centre of the particle, allowing accurate classification of individual particles. We thus could shed light on the inherent conformational landscape of the hypomodified ribosomal RNA within 50S subunits that are capable of assembling into 70S ribosomes. Within the existing structural ensemble, we observe the stable conformations that are sufficiently populated to guide focused classification.

Both types of hypo-modified ribosomes catalyze peptide bond formation at a rate that is three times slower than that of WT ribosomes (Figs. 2, 3). The determined thermodynamic parameters of the PTase reaction revealed that hypo-modified ribosomes have lower activation energy but significantly higher entropic penalty than WT ribosomes. This and their temperature sensitivity (Fig. 3) suggests that hypo-modified ribosomes have a structure less efficient for catalysis. Structures of Δ9 and Δ10 ribosomes indeed show severe misfolding of the PTC (Figs. 5, 7 and 8) with the latter showing the largest heterogeneity (Supplementary Figs. S5, S9, S11–S13).

In particular three MNs in PTCγ seem essential to maintain the native PTC structure. ho^5^C2501, m^2^A2503, and Ψ2504 form a mesh of native tertiary interactions (Supplementary Fig. S8A). When they in the hypo-modified ribosomes adopt non-native conformations, this can result in wide-spread misfolding of the surrounding structure, to a large degree stabilized by alternative stacking interactions (Supplementary Fig. S8B). Interestingly, also the ΔRlmE ribosomes, which contain all MNs except Um2552, contain one class (class 2, Supplementary Fig. S9) with partial disorder of PTCγ, indicating that some alternative conformations are sampled also in WT ribosomes.

During translation, the peptidyl transfer reaction requires the binding and correct orientation of two substrates: a peptidyl-tRNA in the P site and an aminoacyl-tRNA in the A site. Multiple non-active states could be identified, where nucleotides important for coordinating the tRNA substrates are engaged in non-native contacts (Figs 4, 5). Misfolding of the PTC region corroborates the altered activation parameters of the PTase reaction. The acceleration of peptide bond formation on ribosomes is achieved by lowering the entropy of activation (41, 45). It appears that the modifications around PTC play an important role in lowering the entropic penalty of the PTase reaction (Table 1). Both altered activation parameters of the PTase reaction and temperature sensitivity of hypo-modified ribosomes (Fig. 3) agree with the structural importance of modified nucleotides in organizing functional PTC.

Comparison of the native and misfolded PTC structures provides insights into the structural roles of many of the modifications. Critical polar contacts between the ho^5^C2501 base and PTCβ, in particular D2449 (Fig. 6) stabilize the native structure. In absence of the modification, this base can move to an alternative position (Fig. 5C, D). Increased stacking propensity due to base-methylation favours the native intercalation of m^2^A2503 in the PTCα base-stack, but in Δ9 and Δ10 ribosomes it is frequently observed outside this stack (Fig. 5D, Supplementary Fig. S9). The extra polar moiety of the base in pseudouridin compared to the uracil base, locks Ψ2504 in place through solvent-bridged contacts to the backbone (Supplementary Fig. S6). In H89, another important pseudouridin modification, Ψ2457 is positioned by a water-bridged contact with the backbone phosphate (Supplementary Fig. S6). This restricts the movement of the base paired G2494 when U2491 is switched to its IN position (Fig. 8C, D).

The two universally conserved LSU rRNA methylations reside in the A- and P-loops. In the P-loop, the 2’O methylation of G2251 prevents formation of a hydrogen bond to a misfolded conformation of D2449 (Fig. 6C). In the A-loop, the 2’O methyl group of U2552 instead stabilizes the native loop conformation through hydrophobic interactions with the surrounding bases. In its absence, the loop can unfold (Fig. 4E) and disturb nearby nucleotides in PTCδ. Unfolding disables G2553 and U2554 from binding to C75 and C74 of tRNA. Still, both Δ9 and Δ10 ribosomes show very similar PTase activity in the puromycin reaction. This is not surprising, since puromycin acts as an analog of A76 of aatRNA, and is expected to interact mainly with G2583 rather than the A-loop.

Our structures highlight the inherent propensity of large RNAs to adopt alternative stacked conformations. Much like a molecular game of musical chairs, the alternative stacking of one base often displaces another, triggering conformational shifts that propagate across numerous nucleotides (Figs. 5 and 7). We cannot exclude that some of these alternative conformations may occur at low frequency also in WT ribosomes. Evolution has introduced redundancy in rRNA modifications, resulting in the weak phenotypes observed in single knockouts, and where each modification subtly influences the equilibrium between the native structure and a spectrum of potential misfolded states.

## ACKNOWLEDGEMENTS

We acknowledge the use of the Cryo-EM Uppsala facility for grid preparation and screening, funded by the Department of Cell and Molecular Biology, the Disciplinary Domains of Science and Technology and of Medicine and Pharmacy at Uppsala University. Cryo-EM data were collected at the Cryo-EM Swedish National Facility funded by the Knut and Alice Wallenberg Foundation, the Erling-Persson Family Foundation, and the Kempe Foundation; SciLifeLab; Stockholm University; and Umeå University. The data processing was enabled by the Kebnekaise resources at HPC2N provided by the Swedish National Infrastructure for Computing (SNIC) and the Berzelius resource provided by the Knut and Alice Wallenberg Foundation at the National Supercomputer Centre.

## AUTHOR CONTRIBUTIONS

A.L. and R.E. prepared ribosomes and ribosome complexes, A.L. performed biochemistry experiments, A.L. and J.R. analyzed data. D.S.D.L. prepared cryo-EM samples, processed cryo-EM data and modeled and analyzed structures. D.S.D.L., J.R. and M.S. wrote the manuscript with help from all the authors. J.R. and M.S. conceived the study and secured funding.

## DATA AVAILABILITY

The cryo-EM maps and models in this study have been deposited in the Electron Microscopy Data Bank under the accession codes EMDB-55349, 55427–55430, 55432–55437, 55415, 55467–55476, 55618, 55702–55711 and in the Protein Data Bank under the accession codes PDB 9SYH, 9T19–9T1C, 9T1E–9T1J, 9T0Y, 9T2J–9T2S, 9T6M, 9T8K–9T8T. The raw cryo-EM movies have been deposited in the EMPIAR database under the accession code EMPIAR-13296.

