## Supplementary material for "23S rRNA modifications stimulate catalytic activity and prevent the formation of alternative structures"

**Content:**

**Supplementary Tables S1-S5**
**Supplementary Figures S1-S13**
**Supplementary references**

**Supplementary Table S1.** Modified nucleosides of 23S rRNA absent in the *E. coli* strains *Δ9* and *Δ10* and the corresponding modification enzymes**.**

| 23S rRNA position | Modification | Enzyme |
|---|---|---|
| 955 | Ψ | RluC |
| 2069 | $m^7G$ | RlmKL (RlmK domain) |
| 2251* (P-loop) | Gm | RlmB |
| 2445 | $m^2G$ | RlmKL (RlmL domain) |
| 2457* | Ψ | RluE |
| 2498 | Cm | RlmM |
| 2501* | $ho^5C$ | RlhA |
| 2503 | $m^2A$ | RlmN |
| 2504 | Ψ | RluC |
| 2552* (A-loop) | Um | RlmE** |
| 2580 | Ψ | RluC |
| 2604 | Ψ | RluF |
| 2605 | Ψ | RluB |

*Evolutionarily conserved modification site from bacteria to humans

** Present in *Δ10* strain.

**Supplementary Table S2.** Temperature dependence of the rate ($k_{obs}$) of the Tripeptide assay (fMetPhe-tRNA + Pmn)

| | *Δ10* | | *Δ9* | | WT | |
|---|---|---|---|---|---|---|
| Temp °C | $k_{obs}$ sek$^{-1}$ | std | $k_{obs}$ sek$^{-1}$ | std | $k_{obs}$ sek$^{-1}$ | std |
| 15 | 1.83 | 0.38 | 3.08 | 0.32 | 5.23 | 0.39 |
| 20 | 2.28 | 0.55 | 3.85 | 0.30 | 7.85 | 1.00 |
| 25 | 5.45 | 0.93 | 5.50 | 0.45 | 9.91 | 1.27 |
| 30 | 6.50 | 0.73 | 6.90 | 0.41 | 16.01 | 2.18 |
| 37 | 7.49 | 0.93 | 8.42 | 1.03 | 25.89 | 3.02 |

**Supplementary Table S3.** Regions of interest in 23S rRNA

| | Chain | Residues | Structure element[#] |
|---|---|---|---|
| PTCα | 23S | 2058–2063 | H74 |
| PTCβ | 23S | 2447–2453 | H74 |
| PTCɣ | 23S | 2499–2507 | H89, H90 |
| PTCδ | 23S | 2581–2587 | H90 |
| PTCε | 23S | 2607–2610 | H93 |
| P-loop | 23S | 2249–2255 | H80 |
| A-loop | 23S | 2552–2556 | H92 |

[#] Based on (1)

**Supplementary Table S4.** Class distribution after focused 3D classification of the tRNA sites.

| | **APE** | **AP** | **PE** | **E** | **Total A-site** | **Total P-site** |
|---|---|---|---|---|---|---|
| **ΔRlmE** | 52.6% | 9.0% | 24.0% | 14.4% | 61.6% | 85.6% |
| **Δ9** | 50.2% | – | 34.9% | 14.2% | 50.2% | 85.1% |
| **Δ10** | 26.2% | – | 49.6% | 24.1% | 26.2% | 75.8% |

**Supplementary Table S5** Cryo-EM data collection, refinement and validation statistics

| | Δ9 consensus (EMDB-55415) (PDB 9T0Y) | Δ9 class 0 (EMDB-55467) (PDB 9T2J) | Δ9 class 1 (EMDB-55468) (PDB 9T2K) | Δ9 class 2 (EMDB-55469) (PDB 9T2L) | Δ9 class 3 (EMDB-55470) (PDB 9T2M) | Δ9 class 4 (EMDB-55471) (PDB 9T2N) | Δ9 class 5 (EMDB-55472) (PDB 9T2O) | Δ9 class 6 (EMDB-55473) (PDB 9T2P) | Δ9 class 7 (EMDB-55474) (PDB 9T2Q) | Δ9 class 8 (EMDB-55475) (PDB 9T2R) | Δ9 class 9 (EMDB-55476) (PDB 9T2S) |
|---|---|---|---|---|---|---|---|---|---|---|---|
| **Data collection and processing** | | | | | | | | | | | |
| Magnification (kx) | 165 | 165 | 165 | 165 | 165 | 165 | 165 | 165 | 165 | 165 | 165 |
| Voltage (kV) | 300 | 300 | 300 | 300 | 300 | 300 | 300 | 300 | 300 | 300 | 300 |
| Electron exposure (e–/Å$^2$) | 30.0 | 30.0 | 30.0 | 30.0 | 30.0 | 30.0 | 30.0 | 30.0 | 30.0 | 30.0 | 30.0 |
| Defocus range (µm) | -0.6/-1.4 | -0.6/-1.4 | -0.6/-1.4 | -0.6/-1.4 | -0.6/-1.4 | -0.6/-1.4 | -0.6/-1.4 | -0.6/-1.4 | -0.6/-1.4 | -0.6/-1.4 | -0.6/-1.4 |
| Pixel size (Å) | 0.8215 | 0.8215 | 0.8215 | 0.8215 | 0.8215 | 0.8215 | 0.8215 | 0.8215 | 0.8215 | 0.8215 | 0.8215 |
| Symmetry imposed | C1 | C1 | C1 | C1 | C1 | C1 | C1 | C1 | C1 | C1 | C1 |
| Initial particle images (no.) | 1,046,157 | 1,046,157 | 1,046,157 | 1,046,157 | 1,046,157 | 1,046,157 | 1,046,157 | 1,046,157 | 1,046,157 | 1,046,157 | 1,046,157 |
| Final particle images (no.) | 910,722 | 107,941 | 100,949 | 93,493 | 88,813 | 88,625 | 88,358 | 87,602 | 87,185 | 84,723 | 83,033 |
| Map resolution (Å) | 1.87 | 2.02 | 2.06 | 2.07 | 2.06 | 2.06 | 2.07 | 2.08 | 2.08 | 2.22 | 2.07 |
| FSC threshold | 0.143 | 0.143 | 0.143 | 0.143 | 0.143 | 0.143 | 0.143 | 0.143 | 0.143 | 0.143 | 0.143 |
| Map sharpening *B* factor (Å$^2$) | -30.50 | -35.90 | -38.50 | -38.20 | -37.10 | -37.50 | -36.80 | -37.80 | -38.40 | -39.50 | -37.40 |
| **Refinement** | | | | | | | | | | | |
| Initial model(s) used | 9SYH | 9T0Y | 9T0Y | 9T0Y | 9T0Y | 9T0Y | 9T0Y | 9T0Y | 9T0Y | 9T0Y | 9T0Y |
| Model resolution (Å) | 1.85 | 2.05 | 2.07 | 2.08 | 2.08 | 2.08 | 2.08 | 2.09 | 2.09 | 2.24 | 2.09 |
| FSC threshold | 0.5 | 0.5 | 0.5 | 0.5 | 0.5 | 0.5 | 0.5 | 0.5 | 0.5 | 0.5 | 0.5 |
| Model composition | | | | | | | | | | | |
| Non-hydrogen atoms | 151,117 | 14,602 | 15,318 | 15,318 | 15,018 | 15,196 | 14,939 | 15,164 | 15,318 | 15,019 | 15,316 |
| Protein residues | 5,553 | 204 | 212 | 212 | 206 | 197 | 196 | 195 | 212 | 206 | 212 |
| RNA residues | 4,731 | 600 | 631 | 631 | 619 | 631 | 619 | 630 | 631 | 619 | 631 |
| *B* factors (Å$^2$) | 83.37 | 49.73 | 52.06 | 53.29 | 53.05 | 56.15 | 55.46 | 56.73 | 53.55 | 33.44 | 57.02 |
| Protein | 84.21 | 57.82 | 56.50 | 52.92 | 56.34 | 52.20 | 52.72 | 51.79 | 54.88 | 38.07 | 60.41 |
| RNA | 84.79 | 48.42 | 51.30 | 53.14 | 52.41 | 56.39 | 55.54 | 57.08 | 53.17 | 32.62 | 56.41 |
| R.m.s. deviations | | | | | | | | | | | |
| Bond lengths (Å) | 0.0101 | 0.0087 | 0.0089 | 0.0088 | 0.0088 | 0.0088 | 0.0088 | 0.0088 | 0.0088 | 0.0089 | 0.0089 |
| Bond angles (°) | 1.75 | 1.72 | 1.71 | 1.71 | 1.72 | 1.71 | 1.72 | 1.71 | 1.70 | 1.72 | 1.71 |
| Validation | | | | | | | | | | | |
| MolProbity score | 0.93 | 1.31 | 1.21 | 1.29 | 1.33 | 1.35 | 1.33 | 1.27 | 1.27 | 1.31 | 1.28 |
| Clashscore | 0.90 | 1.84 | 1.59 | 1.84 | 2.31 | 2.62 | 2.11 | 1.99 | 1.72 | 2.14 | 1.76 |
| Poor rotamers (%) | 0.07 | 0.65 | 0.00 | 0.00 | 0.00 | 0.00 | 0.68 | 0.00 | 0.00 | 0.00 | 0.00 |
| Ramachandran plot | | | | | | | | | | | |
| Favored (%) | 97.11 | 94.77 | 95.56 | 95.00 | 95.40 | 95.71 | 95.06 | 95.65 | 95.00 | 95.40 | 95.00 |
| Disallowed (%) | 0.09 | 0.00 | 0.00 | 0.00 | 0.00 | 0.00 | 0.00 | 0.00 | 0.00 | 0.00 | 0.00 |

**Supplementary Table 5** Cryo-EM data collection, refinement and validation statistics (Continued)

| | Δ10 consensus (EMDB-55618) (PDB 9T6M) | Δ10 class 0 (EMDB-55702) (PDB 9T8K) | Δ10 class 1 (EMDB-55703) (PDB 9T8L) | Δ10 class 2 (EMDB-55704) (PDB 9T8M) | Δ10 class 3 (EMDB-55705) (PDB 9T8N) | Δ10 class 4 (EMDB-55706) (PDB 9T8O) | Δ10 class 5 (EMDB-55707) (PDB 9T8P) | Δ10 class 6 (EMDB-55708) (PDB 9T8Q) | Δ10 class 7 (EMDB-55709) (PDB 9T8R) | Δ10 class 8 (EMDB-55710) (PDB 9T8S) | Δ10 class 9 (EMDB-55711) (PDB 9T8T) |
|---|---|---|---|---|---|---|---|---|---|---|---|
| **Data collection and processing** | | | | | | | | | | | |
| Magnification (kx) | 165 | 165 | 165 | 165 | 165 | 165 | 165 | 165 | 165 | 165 | 165 |
| Voltage (kV) | 300 | 300 | 300 | 300 | 300 | 300 | 300 | 300 | 300 | 300 | 300 |
| Electron exposure (e–/Å$^2$) | 30.0 | 30.0 | 30.0 | 30.0 | 30.0 | 30.0 | 30.0 | 30.0 | 30.0 | 30.0 | 30.0 |
| Defocus range (μm) | -0.6/-1.4 | -0.6/-1.4 | -0.6/-1.4 | -0.6/-1.4 | -0.6/-1.4 | -0.6/-1.4 | -0.6/-1.4 | -0.6/-1.4 | -0.6/-1.4 | -0.6/-1.4 | -0.6/-1.4 |
| Pixel size (Å) | 0.8215 | 0.8215 | 0.8215 | 0.8215 | 0.8215 | 0.8215 | 0.8215 | 0.8215 | 0.8215 | 0.8215 | 0.8215 |
| Symmetry imposed | C1 | C1 | C1 | C1 | C1 | C1 | C1 | C1 | C1 | C1 | C1 |
| Initial particle images (no.) | 863,376 | 863,376 | 863,376 | 863,376 | 863,376 | 863,376 | 863,376 | 863,376 | 863,376 | 863,376 | 863,376 |
| Final particle images (no.) | 582,240 | 61,890 | 59,988 | 59,781 | 58,711 | 58,616 | 57,630 | 57,497 | 56,658 | 56,208 | 55,261 |
| Map resolution (Å) | 1.87 | 2.18 | 2.21 | 2.20 | 2.22 | 2.19 | 2.22 | 2.23 | 2.22 | 2.22 | 2.23 |
| FSC threshold | 0.143 | 0.143 | 0.143 | 0.143 | 0.143 | 0.143 | 0.143 | 0.143 | 0.143 | 0.143 | 0.143 |
| Map sharpening *B* factor (Å$^2$) | -25.24 | -38.30 | -39.20 | -38.90 | -39.90 | -38.50 | -39.60 | -39.90 | -39.40 | -39.80 | -39.90 |
| **Refinement** | | | | | | | | | | | |
| Initial model(s) used | 9T0Y | 9T6M | 9T6M | 9T6M | 9T6M | 9T6M | 9T6M | 9T6M | 9T6M | 9T6M | 9T6M |
| Model resolution (Å) | 1.85 | 2.22 | 2.25 | 2.23 | 2.25 | 2.25 | 2.26 | 2.26 | 2.25 | 2.25 | 2.26 |
| FSC threshold | 0.5 | 0.5 | 0.5 | 0.5 | 0.5 | 0.5 | 0.5 | 0.5 | 0.5 | 0.5 | 0.5 |
| Model composition | | | | | | | | | | | |
| Non-hydrogen atoms | 150,408 | 14,213 | 14,339 | 14,222 | 15,049 | 14,255 | 14,684 | 14,741 | 14,575 | 14,706 | 14,937 |
| Protein residues | 5,553 | 184 | 193 | 187 | 203 | 196 | 196 | 195 | 193 | 197 | 203 |
| RNA residues | 4,724 | 594 | 596 | 593 | 626 | 591 | 611 | 614 | 607 | 612 | 621 |
| *B* factors (Å$^2$) | 87.77 | 52.86 | 26.19 | 25.68 | 61.18 | 52.79 | 58.01 | 30.94 | 29.65 | 58.42 | 60.59 |
| Protein | 91.40 | 55.66 | 33.14 | 29.28 | 62.60 | 61.26 | 60.23 | 34.32 | 33.10 | 61.92 | 62.93 |
| RNA | 88.26 | 52.38 | 25.21 | 25.09 | 60.91 | 51.59 | 57.63 | 30.42 | 29.12 | 57.88 | 60.23 |
| R.m.s. deviations | | | | | | | | | | | |
| Bond lengths (Å) | 0.0102 | 0.0089 | 0.0089 | 0.0089 | 0.0089 | 0.0089 | 0.0089 | 0.0090 | 0.0089 | 0.0089 | 0.0090 |
| Bond angles (°) | 1.78 | 1.73 | 1.73 | 1.73 | 1.74 | 1.74 | 1.72 | 1.73 | 1.73 | 1.73 | 1.74 |
| Validation | | | | | | | | | | | |
| MolProbity score | 0.94 | 1.14 | 1.38 | 1.23 | 1.32 | 1.21 | 1.28 | 1.30 | 1.25 | 1.35 | 1.27 |
| Clashscore | 0.96 | 1.40 | 1.79 | 1.62 | 2.26 | 1.39 | 2.14 | 1.70 | 1.63 | 1.87 | 1.89 |
| Poor rotamers (%) | 0.07 | 0.00 | 1.37 | 0.00 | 0.66 | 0.00 | 0.67 | 0.00 | 0.00 | 0.67 | 0.66 |
| Ramachandran plot | | | | | | | | | | | |
| Favored (%) | 97.11 | 96.05 | 95.09 | 95.48 | 95.38 | 95.18 | 95.78 | 94.55 | 95.09 | 94.01 | 95.38 |
| Disallowed (%) | 0.09 | 0.00 | 0.00 | 0.00 | 0.00 | 0.00 | 0.00 | 0.00 | 0.00 | 0.00 | 0.00 |

**Supplementary Table 5** Cryo-EM data collection, refinement and validation statistics (Continued)

| | ΔRlmE consensus (EMDB-55349) (PDB 9SYH) | ΔRlmE class 0 (EMDB-55427) (PDB 9T19) | ΔRlmE class 1 (EMDB-55428) (PDB 9T1A) | ΔRlmE class 2 (EMDB-55429) (PDB 9T1B) | ΔRlmE class 3 (EMDB-55430) (PDB 9T1C) | ΔRlmE class 4 (EMDB-55432) (PDB 9T1E) | ΔRlmE class 5 (EMDB-55433) (PDB 9T1F) | ΔRlmE class 6 (EMDB-55434) (PDB 9T1G) | ΔRlmE class 7 (EMDB-55435) (PDB 9T1H) | ΔRlmE class 8 (EMDB-55436) (PDB 9T1I) | ΔRlmE class 9 (EMDB-55437) (PDB 9T1J) |
|---|---|---|---|---|---|---|---|---|---|---|---|
| **Data collection and processing** | | | | | | | | | | | |
| Magnification (kx) | 105 | 105 | 105 | 105 | 105 | 105 | 105 | 105 | 105 | 105 | 105 |
| Voltage (kV) | 300 | 300 | 300 | 300 | 300 | 300 | 300 | 300 | 300 | 300 | 300 |
| Electron exposure (e–/Å$^2$) | 40.6 | 40.6 | 40.6 | 40.6 | 40.6 | 40.6 | 40.6 | 40.6 | 40.6 | 40.6 | 40.6 |
| Defocus range (μm) | -0.4/-1.0 | -0.4/-1.0 | -0.4/-1.0 | -0.4/-1.0 | -0.4/-1.0 | -0.4/-1.0 | -0.4/-1.0 | -0.4/-1.0 | -0.4/-1.0 | -0.4/-1.0 | -0.4/-1.0 |
| Pixel size (Å) | 0.8228 | 0.8228 | 0.8228 | 0.8228 | 0.8228 | 0.8228 | 0.8228 | 0.8228 | 0.8228 | 0.8228 | 0.8228 |
| Symmetry imposed | C1 | C1 | C1 | C1 | C1 | C1 | C1 | C1 | C1 | C1 | C1 |
| Initial particle images (no.) | 736,925 | 736,925 | 736,925 | 736,925 | 736,925 | 736,925 | 736,925 | 736,925 | 736,925 | 736,925 | 736,925 |
| Final particle images (no.) | 513,497 | 46,784 | 54,635 | 53,040 | 15,832 | 57,081 | 55,766 | 56,652 | 51,232 | 53,129 | 69,346 |
| Map resolution (Å) | 1.89 | 2.41 | 2.39 | 2.32 | 3.07 | 2.28 | 2.32 | 2.29 | 2.33 | 2.30 | 2.23 |
| FSC threshold | 0.143 | 0.143 | 0.143 | 0.143 | 0.143 | 0.143 | 0.143 | 0.143 | 0.143 | 0.143 | 0.143 |
| Map sharpening *B* factor (Å$^2$) | -44.2 | -39.60 | -39.70 | -40.30 | -15.40 | -38.80 | -41.00 | -40.00 | -39.60 | -40.00 | -38.70 |
| **Refinement** | | | | | | | | | | | |
| Initial model(s) used | 8CF1, 8CGJ, 8CGK, 7K00 | 9SYH | 9SYH | 9SYH | 9SYH | 9SYH | 9SYH | 9SYH | 9SYH | 9SYH | 9SYH |
| Model resolution (Å) | 2.07 | 2.55 | 2.41 | 2.45 | 3.27 | 2.40 | 2.40 | 2.40 | 2.41 | 2.40 | 2.32 |
| FSC threshold | 0.5 | 0.5 | 0.5 | 0.5 | 0.5 | 0.5 | 0.5 | 0.5 | 0.5 | 0.5 | 0.5 |
| Model composition | | | | | | | | | | | |
| Non-hydrogen atoms | 151,069 | 15,294 | 15,298 | 15,254 | 14,819 | 15,297 | 15,295 | 15,038 | 15,256 | 15,041 | 15,298 |
| Protein residues | 5,553 | 211 | 211 | 205 | 203 | 211 | 211 | 211 | 205 | 211 | 211 |
| RNA residues | 4,734 | 629 | 629 | 629 | 617 | 629 | 629 | 617 | 629 | 617 | 629 |
| *B* factors (Å$^2$) | 104.74 | 31.14 | 25.23 | 27.35 | 88.81 | 29.08 | 26.47 | 24.97 | 29.43 | 25.73 | 28.35 |
| Protein | 110.26 | 31.59 | 25.82 | 29.15 | 87.97 | 27.81 | 26.73 | 28.51 | 27.11 | 28.51 | 27.06 |
| RNA | 104.56 | 30.90 | 24.92 | 26.88 | 88.95 | 29.05 | 26.23 | 24.29 | 29.50 | 25.13 | 28.32 |
| R.m.s. deviations | | | | | | | | | | | |
| Bond lengths (Å) | 0.0094 | 0.0094 | 0.0095 | 0.0093 | 0.0098 | 0.0094 | 0.0094 | 0.0093 | 0.0094 | 0.0094 | 0.0094 |
| Bond angles (°) | 1.69 | 1.81 | 1.79 | 1.78 | 1.85 | 1.77 | 1.79 | 1.79 | 1.79 | 1.79 | 1.80 |
| Validation | | | | | | | | | | | |
| MolProbity score | 0.90 | 1.33 | 1.43 | 1.39 | 1.44 | 1.24 | 1.28 | 1.34 | 1.41 | 1.39 | 1.32 |
| Clashscore | 0.82 | 2.73 | 2.22 | 2.48 | 2.59 | 2.01 | 2.01 | 1.92 | 2.19 | 2.30 | 2.01 |
| Poor rotamers (%) | 0.13 | 0.64 | 1.27 | 0.65 | 0.65 | 0.00 | 0.00 | 0.64 | 0.65 | 0.64 | 0.64 |
| Ramachandran plot | | | | | | | | | | | |
| Favored (%) | 97.20 | 96.09 | 94.97 | 94.74 | 94.15 | 96.09 | 95.53 | 94.41 | 93.64 | 94.41 | 94.97 |
| Disallowed (%) | 0.09 | 0.00 | 0.00 | 0.00 | 0.00 | 0.00 | 0.00 | 0.00 | 0.00 | 0.00 | 0.00 |

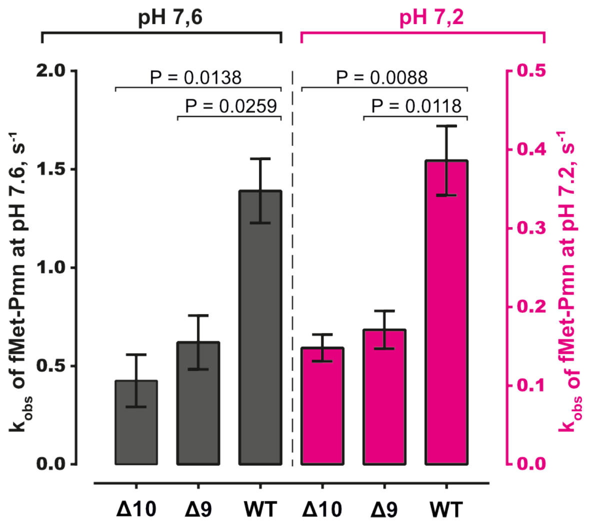


**Supplementary Figure S1.** The effect of pH on the puromycin reaction with fMet-tRNA (dipeptide assay) is similar for WT and hypo-modified ribosomes.

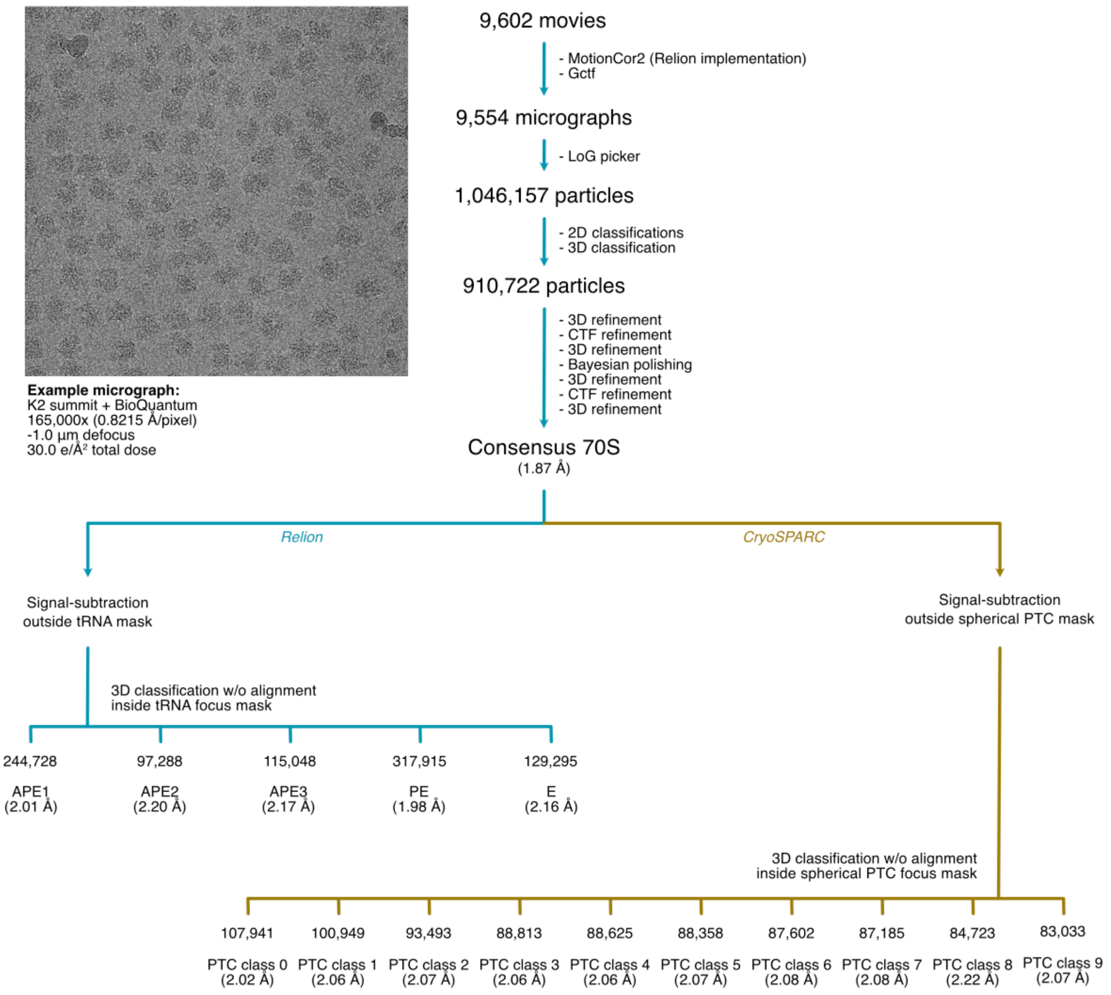


**Supplementary Figure S2.** Cryo-EM single-particle analysis workflow and example micrographs for the Δ9 reconstructions.

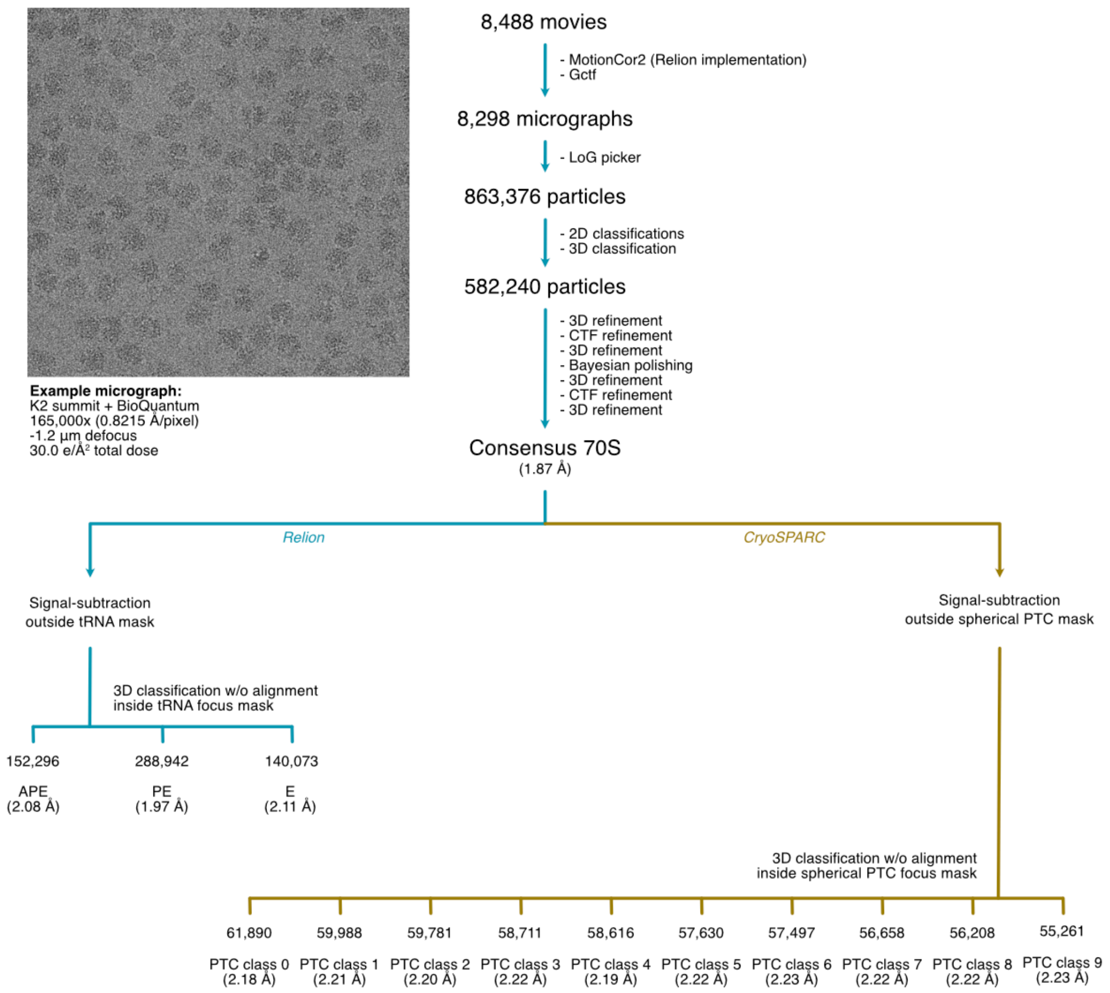


**Supplementary Figure S3.** Cryo-EM single-particle analysis workflow and example micrographs for the Δ10 reconstructions.

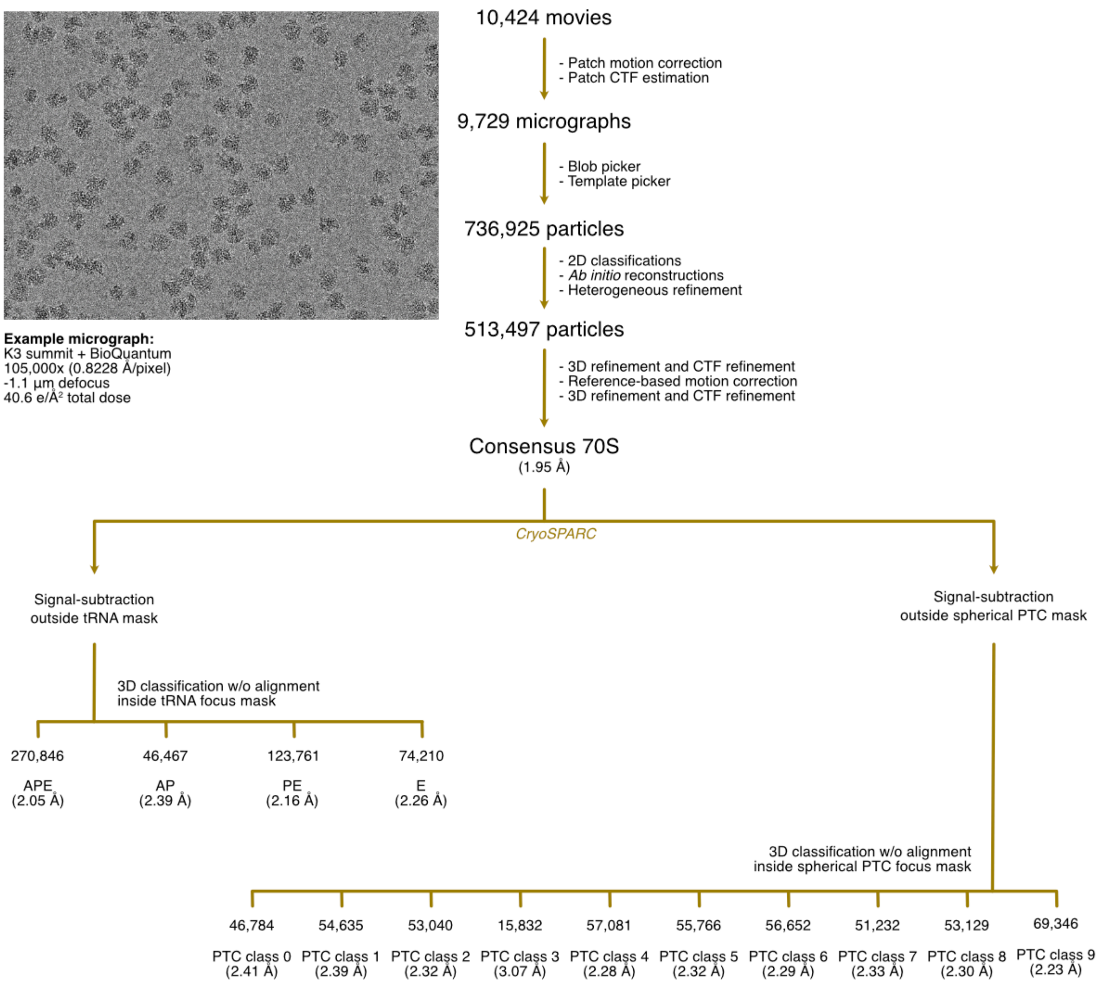


**Supplementary Figure S4.** Cryo-EM single-particle analysis workflow and example micrographs for the ΔRlmE reconstructions.

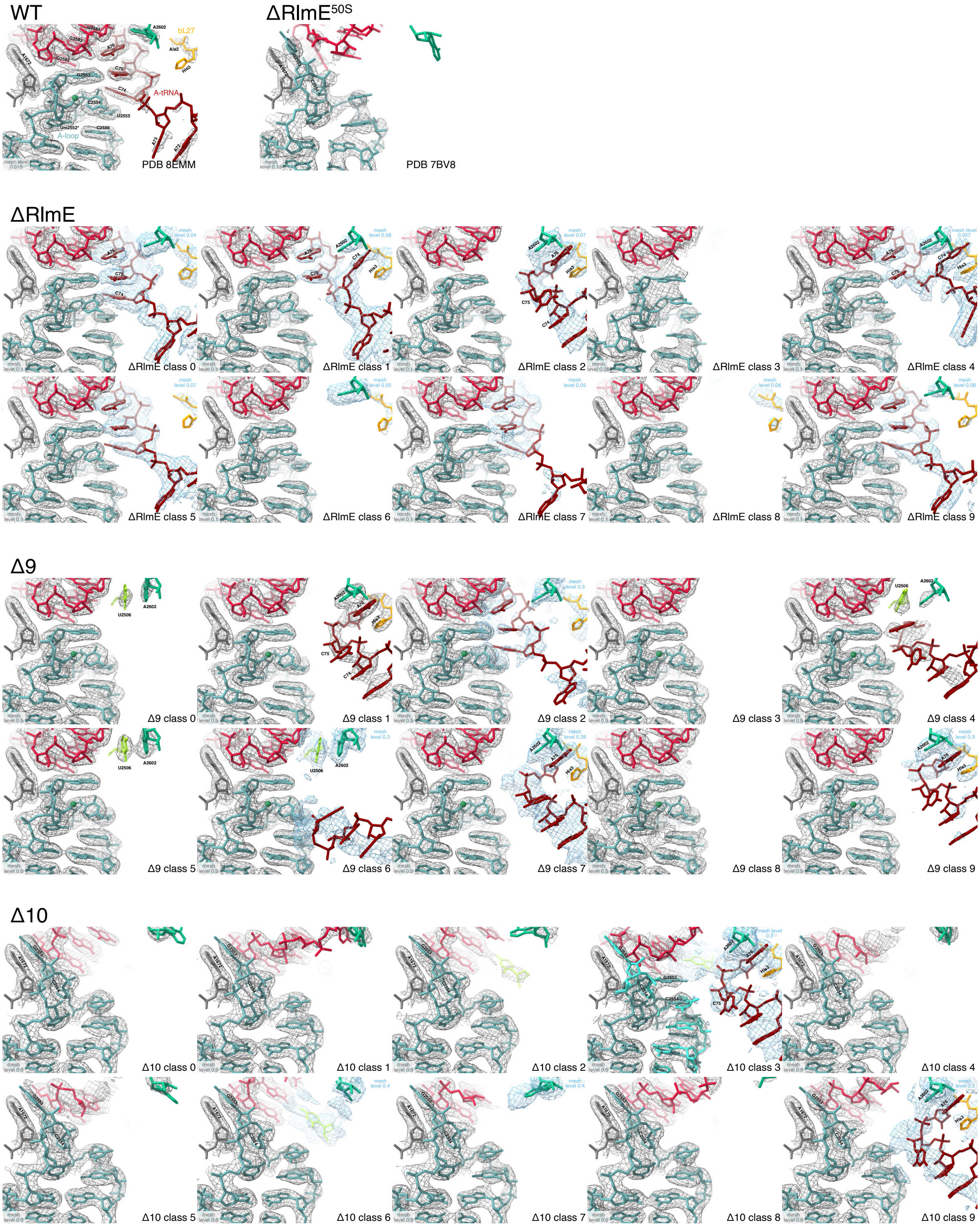


**Supplementary Figure S5.** A-loop conformation for all PTC classes. The view is the same as in Fig. 4A–E. Reference structures are WT 70S with A- and P-site tRNA (PDB 8EMM (2)) and 50S ΔRlmE (PDB 7BV8). Δ10 class 3 has an A-loop (pale blue) with WT conformation, but an alternative conformation is with G2553 and C2554 flipped out (cyan). All other Δ10 classes have G2553 and C2554 flipped out. Δ9 class 2 (but no Δ10 class) has week density for a WT conformation of the CCA-end of the A-site tRNA (maroon). Several classes have density for the A-site tRNA being captured between A2602 (mint green) and His3 of bL27 (orange): ΔRlmE classes 1, 2 and 4; Δ9 classes 1, 7 and 9; Δ10 classes 3 and 9. U2506 (lime green) stack with A2602 (mint green) in many Δ9 classes (0, 4, 5 & 6), blocking A76 binding.

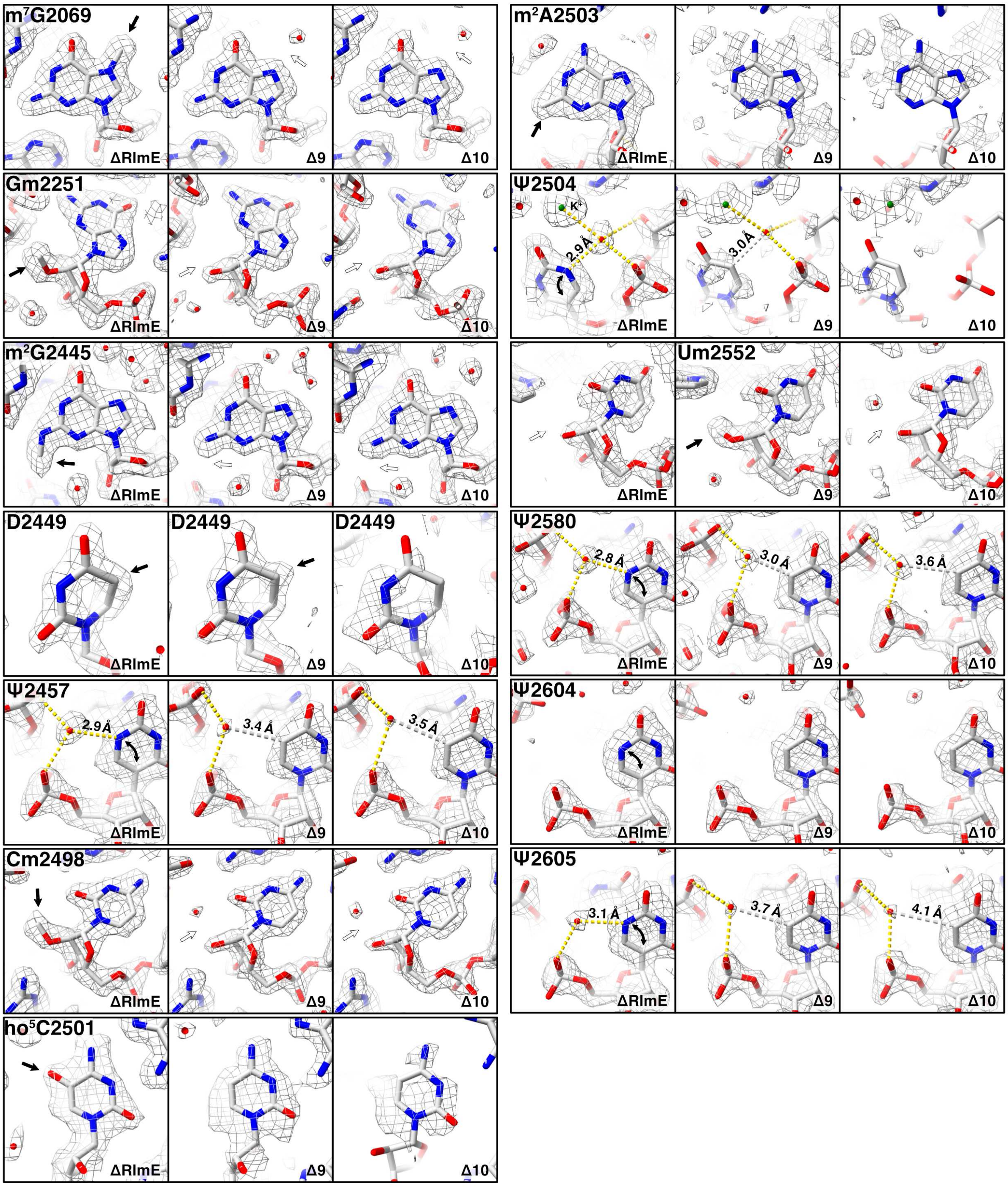


**Supplementary Figure S6.** Map support for post-transcriptional modifications in consensus reconstructions. Black arrows indicate the presence and white arrows the absence of modifications.

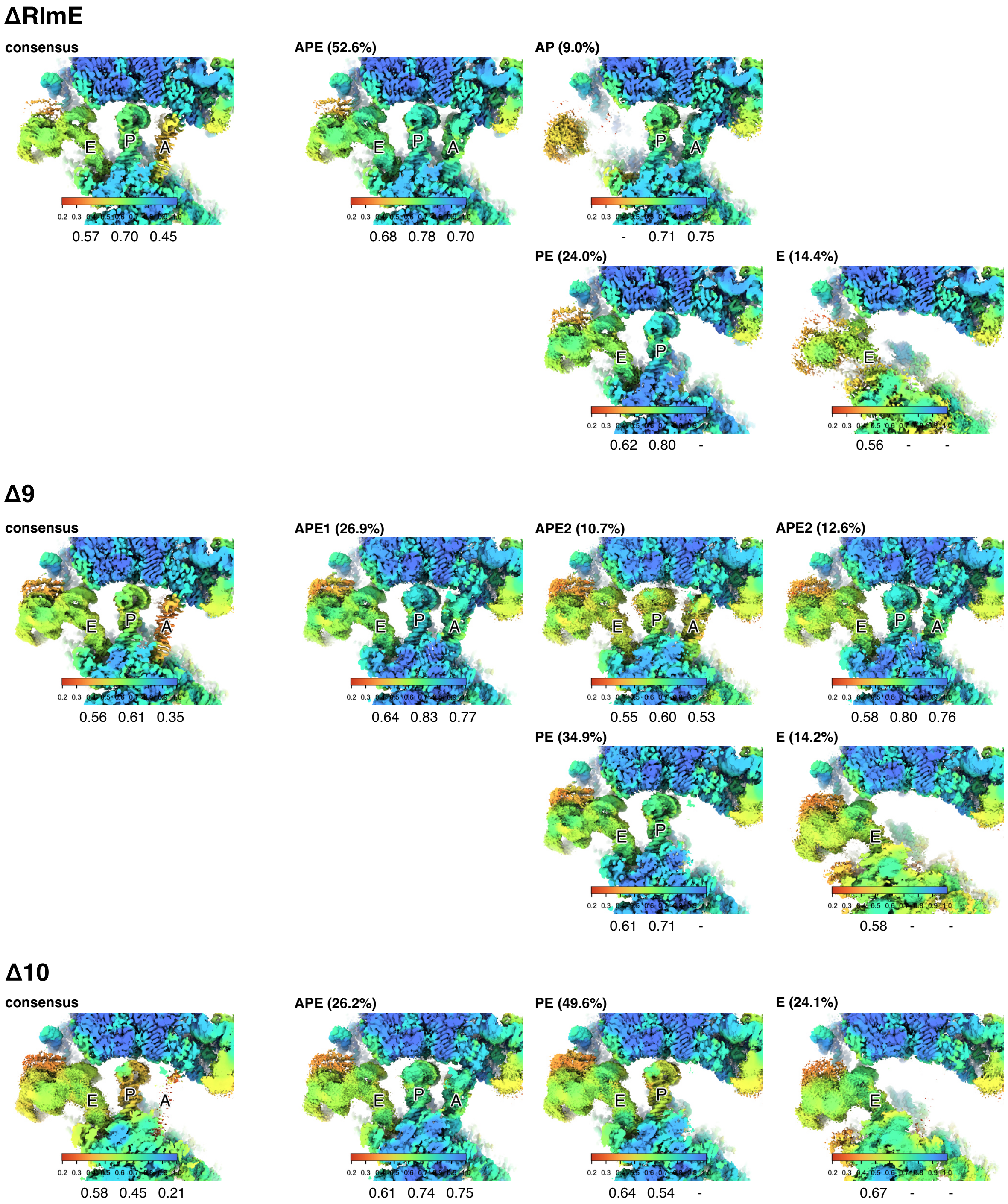


**Supplementary Figure S7.** tRNA occupancy analysis for consensus reconstructions and tRNA classes using Occupy (3). Colors represent the estimated local occupancy. Values below each figure represent the occupancy at the middle of each tRNA molecule. Percentages indicate the fraction of particles in that class.

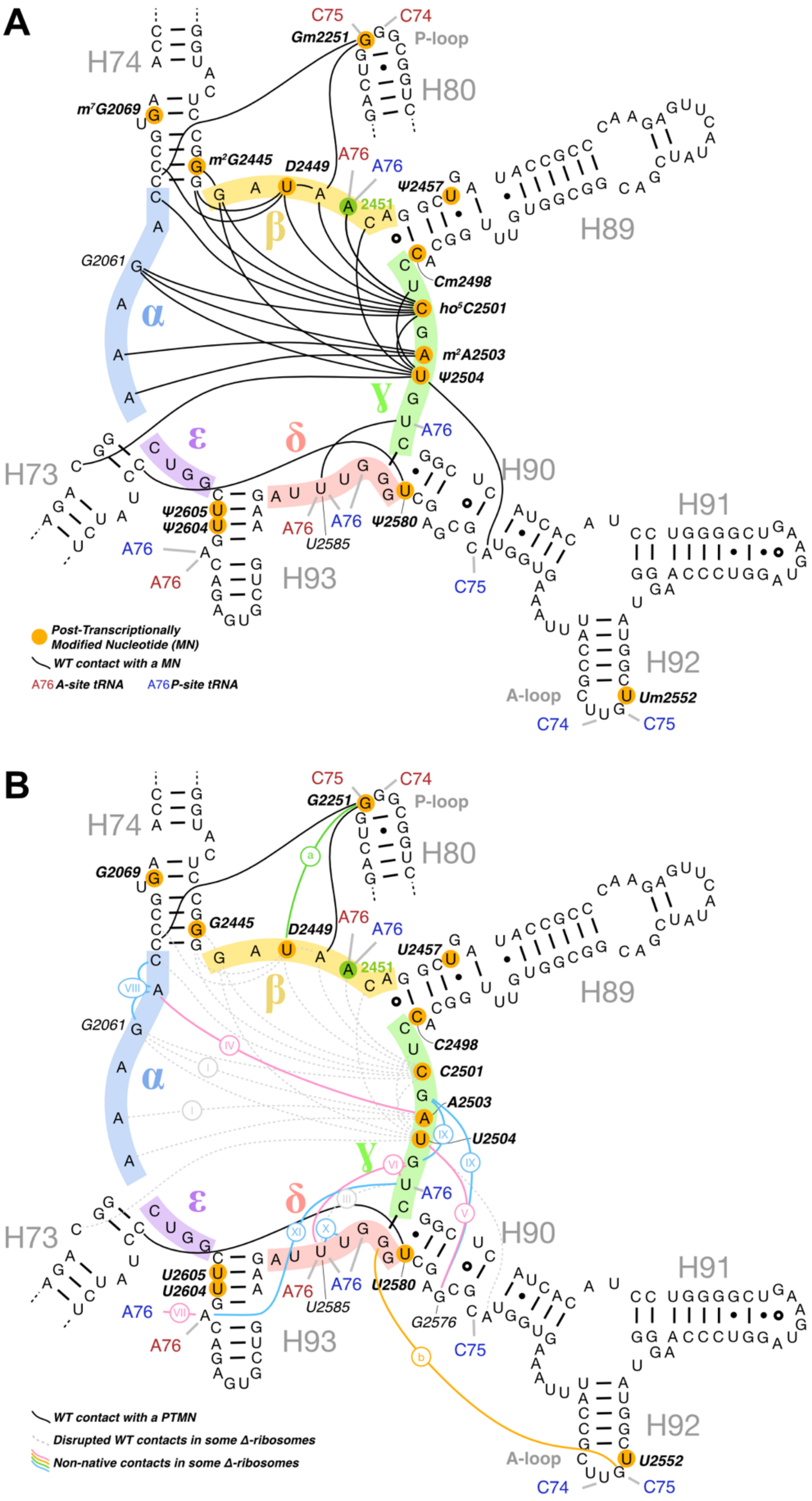


**Supplementary Figure S8.** Tertiary contacts in the PTC region. **A**. Secondary structure diagram as in Fig. 1B with tertiary contacts from post-transcriptionally modified nucleotides in the WT ribosome (PDB 8EMM) indicated with black lines. **B**. Disrupted native contacts (dashed gray lines) and non-native contacts (colors) observed in some D9 and D10 classes. Roman numbers I–XI as shown in Fig. 7. Contact a is shown in Fig. 6C and contact b in Fig. 4F.

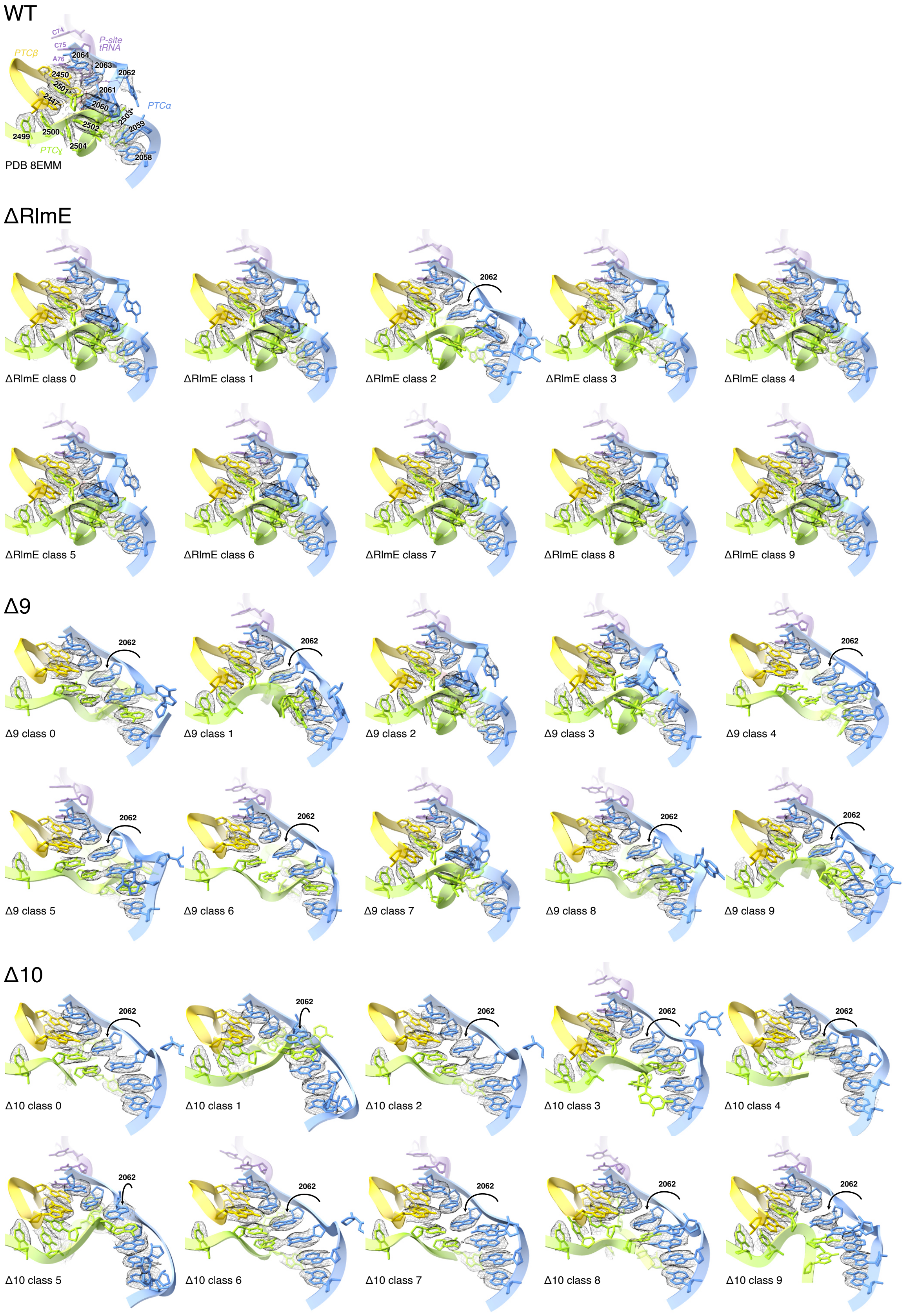


**Supplementary Figure S9.** Misfolding of the PTC-ring. The view and colors are the same as in Fig. 5. Most Δ10 classes have A2062 flipped in. Unexpectedly, ΔRlmE class 2 also shows A2062 flipped in and PTCɣ partially disordered.

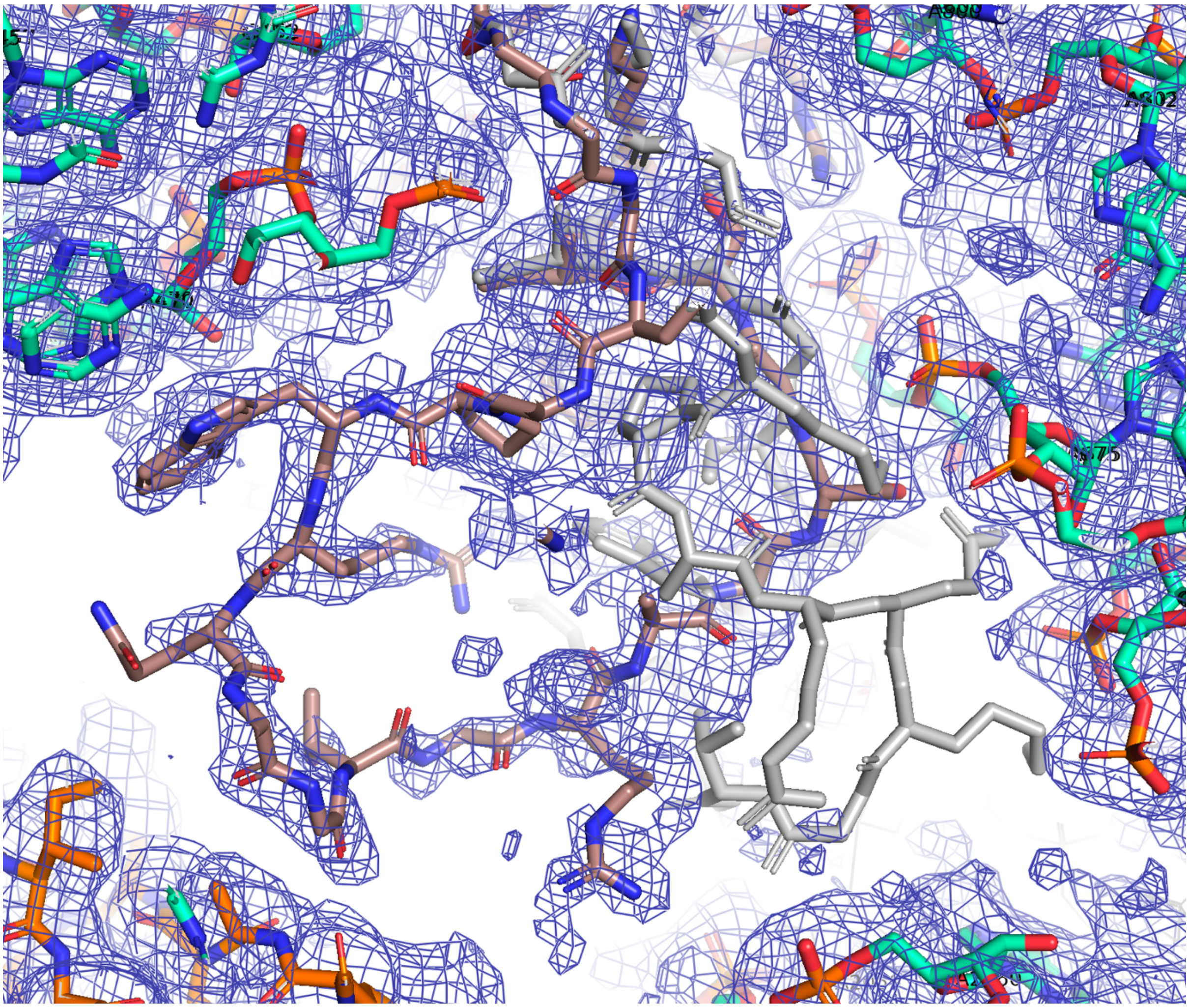


**Supplementary Figure S10.** Alternative folding of loop 55–73 in r-protein uL4. Consensus Δ10 is shown in brown and WT conformation (PDB 8EMM (2)) in grey.

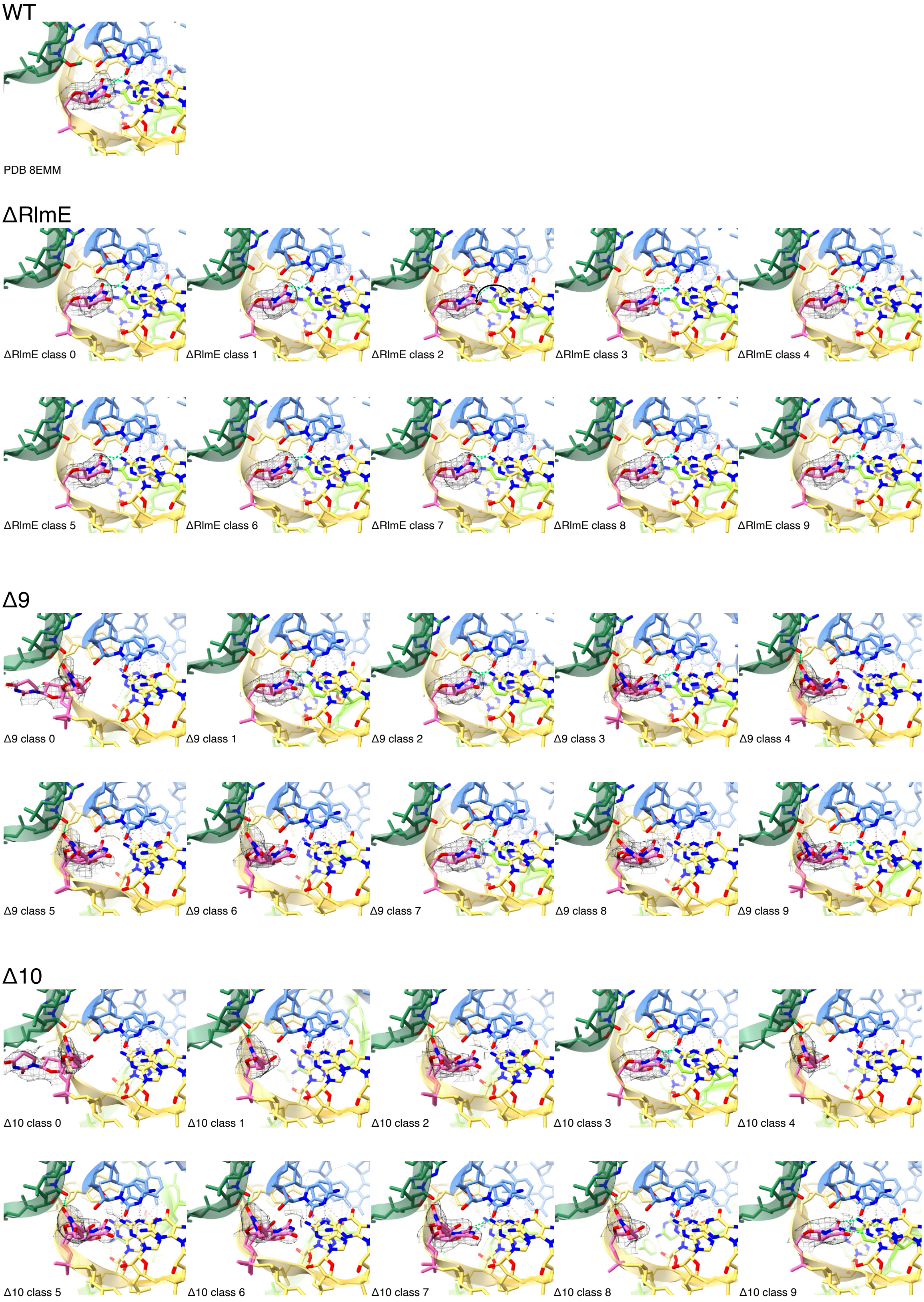


**Supplementary Figure S11.** Conformation of $ho^5$C2501 and D2449. The view and colors are the same as in Fig. 6. Most Δ10 classes (0, 1, 2, 4, 5, 6, 7, 8) have D2449 rotated. Δ10 class 0 also shows yet another alternative conformation for D2449.

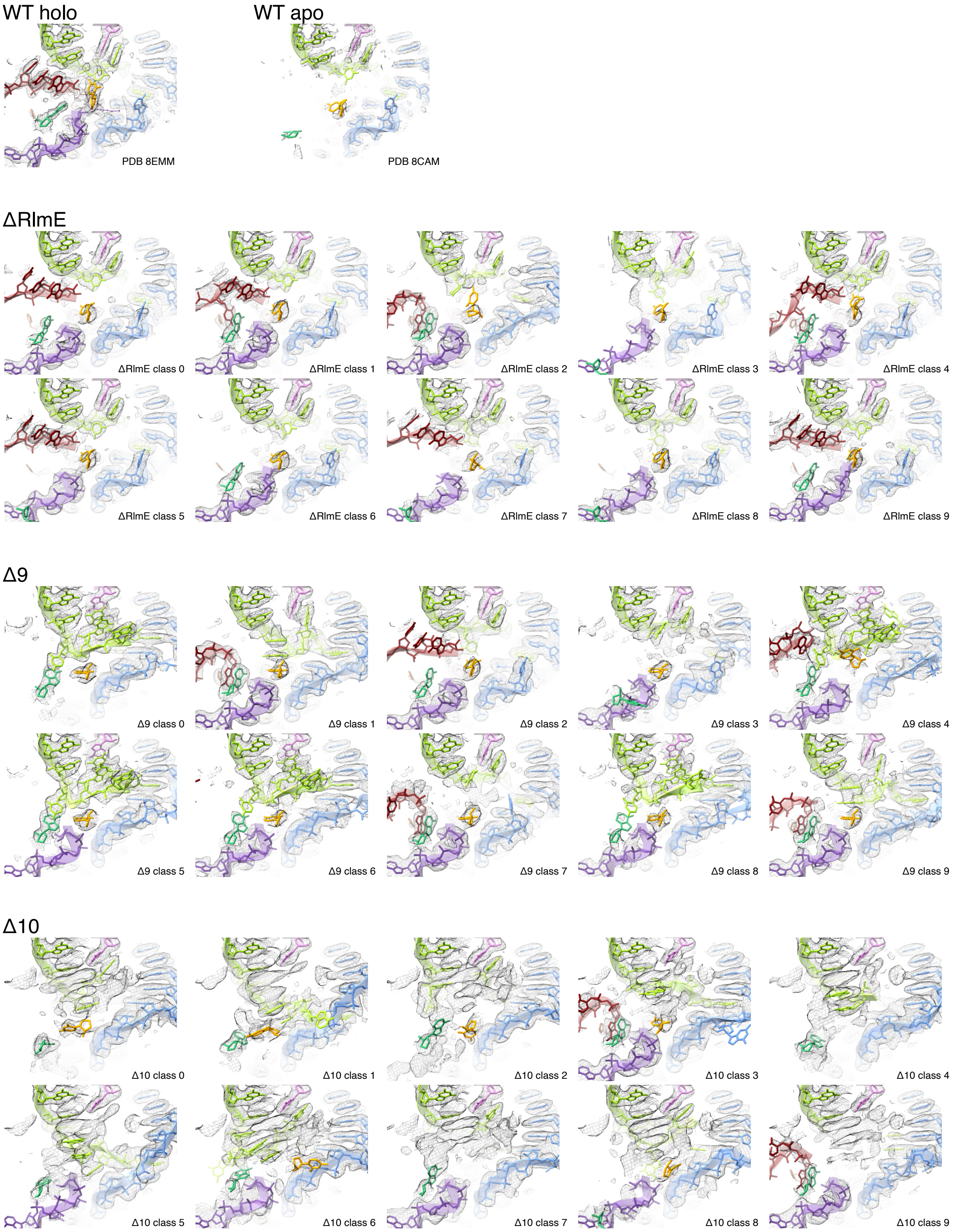


**Supplementary Figure S12.** Alternative stacking at the PTC. The view and colors are the same as in Fig. 7. Reference structures are WT with A- and P-site tRNA (PDB 8EMM (2)) and WT without tRNA (PDB 8CAM). Most Δ10 classes have strong density for stacked bases, but weak density for the backbone of PTCɣ and PTCδ.

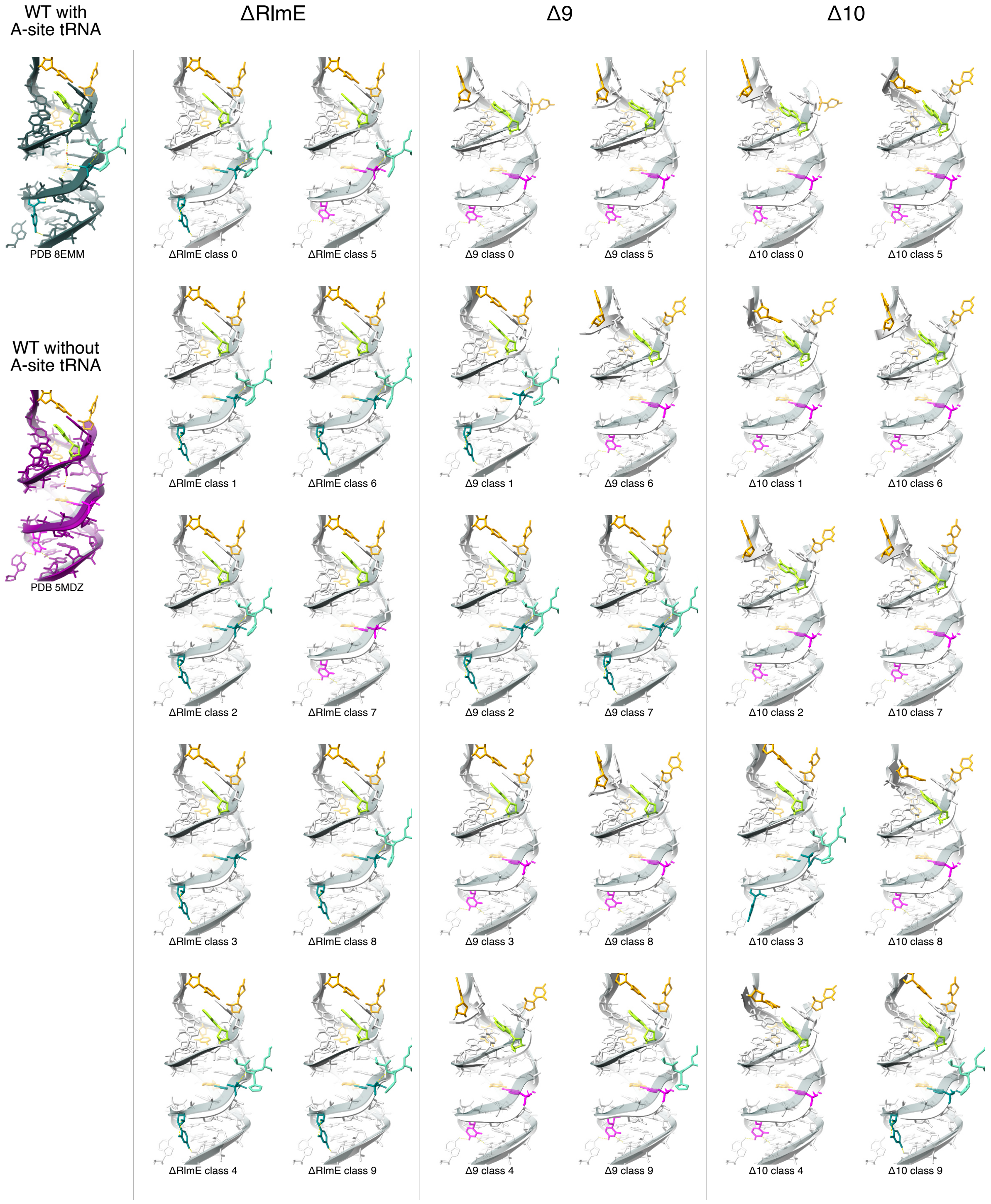


**Supplementary Figure S13.** Switching of U2491 and deformation of H89. The view and colors are the same as in Fig. 8. Reference structures are WT with A- and P-site tRNA (PDB 8EMM (2)) and WT without A-site tRNA (PDB 5MDZ (4)). U2491 in the OUT conformation is shown in teal and in the IN conformation in magenta.
